# Insights from a genome-wide truth set of tandem repeat variation

**DOI:** 10.1101/2023.05.05.539588

**Authors:** Ben Weisburd, Grace Tiao, Heidi L. Rehm

**Affiliations:** Program in Medical and Population Genetics, Broad Institute of MIT and Harvard, Cambridge, MA, USA; Analytic and Translational Genetics Unit, Massachusetts General Hospital, Boston, MA, USA; Center for Genomic Medicine, Massachusetts General Hospital, Boston, MA, USA

## Abstract

Tools for genotyping tandem repeats (TRs) from short read sequencing data have improved significantly over the past decade. Extensive comparisons of these tools to gold standard diagnostic methods like RP-PCR have confirmed their accuracy for tens to hundreds of well-studied loci. However, a scarcity of high-quality orthogonal truth data limited our ability to measure tool accuracy for the millions of other loci throughout the genome. To address this, we developed a TR truth set based on the Synthetic Diploid Benchmark (SynDip). By identifying the subset of insertions and deletions that represent TR expansions or contractions with motifs between 2 and 50 base pairs, we obtained accurate genotypes for 139,795 pure and 6,845 interrupted repeats in a single diploid sample. Our approach did not require running existing genotyping tools on short read or long read sequencing data and provided an alternative, more accurate view of tandem repeat variation. We applied this truth set to compare the strengths and weaknesses of widely-used tools for genotyping TRs, evaluated the completeness of existing genome-wide TR catalogs, and explored the properties of tandem repeat variation throughout the genome. We found that, without filtering, ExpansionHunter had higher accuracy than GangSTR and HipSTR over a wide range of motifs and allele sizes. Also, when errors in allele size occurred, ExpansionHunter tended to overestimate expansion sizes, while GangSTR tended to underestimate them. Additionally, we saw that widely-used TR catalogs miss between 16% and 41% of variant loci in the truth set. These results suggest that genome-wide analyses would benefit from genotyping a larger set of loci as well as further tool development that builds on the strengths of current algorithms. To that end, we developed a new catalog of 2.8 million loci that captures 95% of variant loci in the truth set, and created a modified version of ExpansionHunter that runs 2 to 3x faster than the original while producing the same output.

## Introduction

Tandem repeats (TRs) are sequences consisting of a nucleotide motif that repeats consecutively. They have traditionally been called short tandem repeats (STRs) for motif sizes between 1 and 6 base pairs (bp), and variable number tandem repeats (VNTRs) for larger motifs, but may be collectively referred to as TRs. There are millions of tandem repeat loci scattered throughout the human genome. More than 50 of these are known to cause rare monogenic diseases when they expand beyond a certain number of repeats, termed the pathogenic threshold^1^. The thresholds range from as low as 7 or 8 repeats of a GCG motif at the *PABPN1* locus resulting in oculopharyngodistal myopathy^2^, to as high as 1000 or more repeats of a TTTCA motif at the *YEATS2* locus causing familial adult myoclonic epilepsy^3^. Tandem repeat variation also contributes to a range of other human phenotypes^4–7^.

The first generation of tools for genotyping tandem repeats using short read sequencing data ^8, 9^ only considered reads that overlapped the TR locus and so could only call expansions shorter than read length. Later, ExpansionHunter^10^ and GangSTR^11^ integrated information from mismapped fully-repetitive reads to accurately call allele sizes both longer and shorter than read length. Most recently, ExpansionHunterDenovo^12^ and STRling^13^ demonstrated significant improvements in computational efficiency, albeit with certain new limitations - such as only being able to detect expansions *longer* than read length.

Until recently, rare disease research based on short read sequencing data focused primarily on single nucleotide variants (SNVs) and structural variants (SVs) partly due to skepticism about the ability of short read sequencing data to accurately capture TR variant lengths. However, recent studies involving large rare disease cohorts ^14, 15^ showed that, for known disease-associated loci, genotypes from tools such as ExpansionHunter were highly consistent with gold-standard diagnostic methods. Additionally, multiple groups have begun using tandem repeat genotyping tools for short read data to discover novel monogenic TR loci^16^.

However, there is ambiguity about the best approach to use for genotyping TRs genome-wide in rare disease cohorts. Open questions remain around the set of loci to genotype, the accuracy of existing tools at these loci, and optimal call filtering strategies. To improve our ability to answer these questions, we set out to develop a genome-wide TR truth set.

As our starting point, we chose the Synthetic Diploid Benchmark^17^. This is a unique dataset that uses haplotype-resolved assemblies of PacBio data from two homozygous cell lines to identify all variants in the CHM1-CHM13 synthetic diploid sample (SynDip). Because these variant calls are based on the alignment of entire genome assemblies (rather than individual long reads) to the reference genome, the SynDip genotypes are more reliable than those produced by short read or even ordinary long read pipelines^18^. Additionally, since one of the two homozygous samples that makes up SynDip - CHM13 - is the basis of the telomere-to-telomere (T2T) reference genome, we were able to validate TR genotypes via liftover and comparison to the T2T reference sequence. Below, after describing how we derived and validated the truth set, we evaluate the accuracy of widely-used tools: ExpansionHunter, GangSTR, HipSTR, and ExpansionHunterDenovo. Additionally, we test the completeness of publicly available TR catalogs based on their ability to capture truth set variant loci. To support broader genome-wide studies, we create and share a catalog of 2.8M loci, as well as an optimized version of ExpansionHunter. Finally, because the truth set provides a comprehensive snapshot of TR variants in a diploid human genome, we use it to explore overall properties of simple TRs such as their relative mutation rates and their distribution across genes with different loss-of-function and missense constraint scores.

## Results

### Deriving a TR truth set from the Synthetic Diploid Benchmark

The Synthetic Diploid Benchmark provides genotypes for 5,182,765 SNV, insertion and deletion variants, as well as a set of high-confidence regions spanning 2.71 gigabases where genotypes are highly accurate. We started by selecting all 518,281 insertions and deletions (which we refer to as ins/del variants rather than indels since indels typically only include variant sizes up to 50bp) located within high-confidence regions (genomic intervals where haploid assemblies for CHM1 and CHM13 could be aligned to the reference genome without issues and with sufficient mapping quality^17^) (**Figure 1A**). We hypothesized that a subset of these ins/del variants represented simple TR expansions and contractions that consisted of either pure repeats or repeats with a simple interruption pattern. To identify this subset, we created a filtering algorithm that uses two steps to determine if an ins/del variant represents a TR allele. The first step looks at only the inserted or deleted bases - which we refer to as the variant sequence - and checks whether they consist entirely of repeats of some smaller motif. If not, it considers the motif to be the entire variant sequence. In the second step, the algorithm extends the repeat(s) into the reference context by checking for additional copies of the motif immediately to the left and to the right of the insertion or deletion, as illustrated in **Figures 1B-E**.

**Figure 1:**
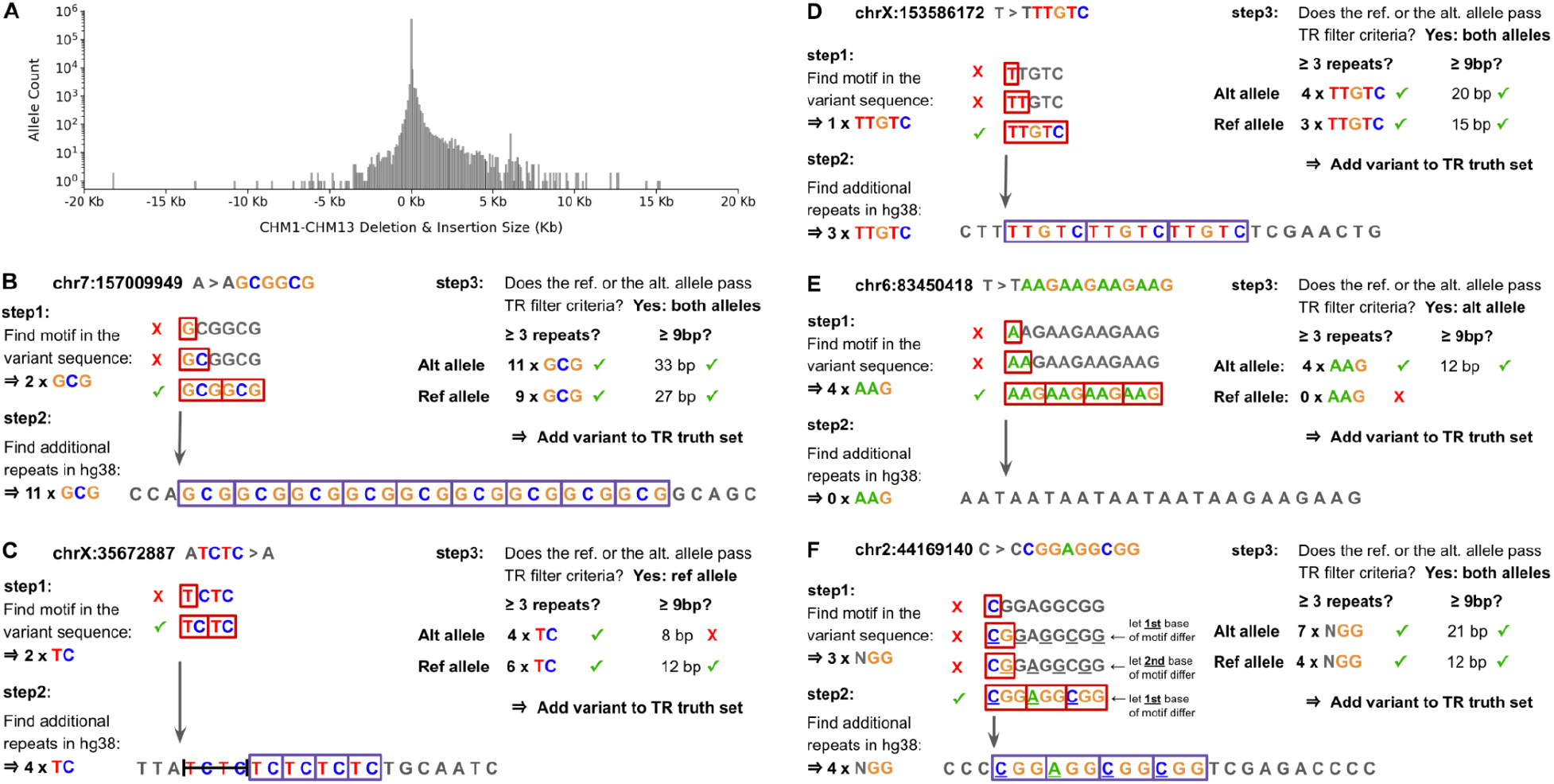
Deriving a TR truth set from the Synthetic Diploid Benchmark. **A. Ins/del allele size distribution in the synthetic diploid benchmark.** A histogram of allele sizes for all 562,891 deletions and insertions found within SynDip high-confidence regions. On the x axis, positive values represent insertion sizes, while negative values represent deletion sizes in kilobases. **B. Example 1: How the TR filter determines if an insertion represents a TR expansion.** To decide whether the A > AGCGGCG 6bp insertion in SynDip at position chr7:157009949 is a pure tandem repeat variant, the filter first uses brute force k-mer search on just the GCGGCG variant sequence to find that it consists of 2 x GCG repeats. Then, in step 2, it looks for additional GCG repeats in the reference immediately to the left and right of the insertion point and finds 9 more repeats there. Since there are 11 total GCG repeats spanning 33 base pairs, this insertion passes the filter criteria (≥3 repeats and spanning ≥9 bp) and is included in the truth set as a TR expansion with a true allele size of 11 x GCG repeats and locus coordinates chr7:157009950-157009976. **C. Example 2: How the TR filter determines if a deletion represents a TR contraction.** To decide whether this 4bp deletion is a pure TR variant, the filter first finds that the deleted bases consist of 2 x TC repeats and then finds 4 additional TC’s in the reference immediately to the right of the deletion. It concludes that the allele is a TR contraction with a true allele size of 4 x TC and locus coordinates chrX:35672888-35672899. **D. Example 3: No repeats in the variant sequence.** When the filter finds that it can’t divide the variant sequence into pure repeats of a smaller motif, it takes the motif to be the entire variant sequence. Then, it checks the reference context and finds 3 more TTGTC’s there. This allele is then added to the truth set as 4 x TTGTC at locus chrX:153586173-153586188 because it has ≥ 3 repeats and spans ≥ 9bp overall. **E. Example 4: No adjacent repeats in the reference context.** Since the filter starts with the variant sequence in step 1, it is able to detect TRs even when they have no matching repeats in their immediate reference context. In this example, there are 4 x AAG repeats in the variant sequence and none immediately to the left or right of the insertion point. The 4 x AAG repeats pass the filter criteria of having ≥ 3 repeats and spanning ≥ 9bp. Alternatively, this example could reasonably be considered an interrupted repeat with an AAN motif that has at least 8 matching repeats in the reference sequence. However, because the filter implementation only tests a variant for interrupted repeats if it doesn’t pass the filtering criteria as a pure repeat, this variant is added to the truth set as a pure non-reference allele with 4 x AAG repeats. **F. Example 5: Detecting TRs that have interruptions.** When the filter determines that an ins/del variant doesn’t pass thresholds as a pure tandem repeat, it retests the sequence, but this time allowing one position to vary across repeats of the motif. In step 1 of this retest phase, it performs a brute force search to find that the variant sequence consists of 3 repeats of an NGG motif where the first position varies across repeats. In step 2, it extends the repeats into the reference context and finds 4 additional NGG repeats. It then adds this allele to the truth set as a 7 x NGG allele at locus chr2:44169141-44169152.

The filter concludes that an ins/del variant is a TR if the repeats in the variant sequence plus the reference context A) collectively span at least 9bp and B) consist of at least 3 repeats of the motif. The minimum threshold of 3 repeats is based in part on some of the known disease-associated loci such as C9orf72 having only 3 repeats in the hg38 reference. Applying the 3 repeat minimum to the total number of repeats rather than only those in the reference sequence enables the filter to detect TR variants with 2, 1, or even 0 repeats in the reference (**Figure 1D**). Also, although we mostly limit our analysis to motifs of size 2 to 50bp, the same filtering approach works for any motif size, including homopolymers.

Initially, the filter only looks for pure repeats. However, if an ins/del variant doesn’t pass the filtering criteria, the algorithm retests the variant while allowing one position within the motif to vary across repeats. This enables the detection of loci with interruptions similar to those seen in 12 known disease-associated loci with GCN motifs where N is variable (**Figure 1F**). For multi-allelic variants, the filtering procedure is applied separately to each alternate allele, and the overall variant passes the filter only if both alternate alleles pass and have the same motif. For every ins/del variant that passes, the filter then outputs all the information needed to run TR genotyping tools on the locus and evaluate results, including the repeat motif, the exact start and end coordinates of repeats it identified in the reference, and the true TR genotype based on the ins/del variant’s genotype within the SynDip variant call file (VCF).

### Validating the TR truth set

The CHM13 cell line serves as the basis of the telomere-to-telomere (T2T) reference genome^19^. This allowed us to validate truth set variants by 1) lifting them over to T2T and 2) checking that, for each variant, at least one allele matched the T2T reference to within +/- 2 repeats. Additionally, to confirm the consistency of the liftover process, we then 3) lifted the variants back over to hg38 and 4) checked that their hg38 position didn’t change after the hg38 ⇒ T2T ⇒ hg38 liftover. When we performed this validation on the 73,361 expansions in the truth set, we found that 68,304 (93.1%) passed all steps. **Table 2** shows the total number of variants that failed each validation step, and **Supplementary Figure 1C** provides additional details about the validation workflow.

**Table 1:** Applying the TR filter to SynDip variants. This table provides a summary of the stages involved in filtering SynDip variants to the subset that represents TR expansions or contractions. Since, at each stage, some fraction of variants is multi-allelic, the “# of alleles” column has a larger number than the “#of variant loci”. The “% of alleles” column represents the fraction of alleles that passes each of the 2 filtering stages.

| Step | Description | # of variant loci | # of alleles | % of alleles |
| --- | --- | --- | --- | --- |
|  | Start with all variants in SynDip high-confidence regions | 4,081,549 | 4,129,306 | 100.0% |
| 1 | Keep only insertions and deletions | 518,281 | 556,825 | 13.5% |
| 2 | Keep only insertions and deletions that represent TR expansions or contractions based on these criteria:<br>• span $\geq 9$ base pairs<br>• have $\geq 3$ repeats<br>• motif size $\geq 2$bp and $\leq 50$bp | 157,601 | 188,730 | 4.6% |

**Table 2:** Validating TR variants against the T2T reference genome. This table provides a brief overview of the TR validation procedure and summarizes the number and percentage of TR variants that failed each of the 4 validation steps. A more detailed flowchart and statistics are provided in **Supplementary Figure 1**.

| Step | Description | # failed | % failed |
| --- | --- | --- | --- |
|  | Start with all 157,601 TR variants in SynDip high-confidence regions |  |  |
| 1 | Liftover TR variants from hg38 ⇒ T2T | 1,626 | 1.0% |
| 2 | Check that at least one allele matches the T2T reference sequence | 9,313 | 5.9% |
| 3 | Lift variants back over from T2T ⇒ hg38 | 22 | 0.0% |
| 4 | Check if the variant's position changed after hg38 ⇒ T2T ⇒ hg38 liftover | 0 | 0.0% |

Due to a technical limitation of existing liftover tools, we could not perform liftover on 10,783 out of 74,888 contractions, as well as 9,203 out of 9,352 mixed multiallelic variants defined as two non-reference alleles where one was a contraction while the other was an expansion. These variants encountered a liftover error because the deletions overlapped boundaries between liftover chain intervals. Since this error is not an indicator of poor variant quality, but is instead a limitation of the liftover algorithm, we added these 19,986 contractions and mixed multiallelic variants to the truth set without validation. Of the 64,254 contractions and mixed multiallelic variants that avoided this error, 58,350 (90.8%) passed all validation steps.

Comparing allele counts, 5,924 out of 95,104 (6.2%) contraction alleles and 5,716 out of 93,626 (6.1%) expansion alleles failed validation. Additionally, alleles that differed from the hg38 reference by more repeats failed at a higher rate than alleles that were closer in size to the hg38 reference allele. However, the size distribution of alleles that failed validation was symmetrical around the reference allele size (**Supp. Figure 1A)**, and furthermore, was approximately uniform across motif sizes (**Supp. Figure 1B)**. Consequently, the validation procedure did not significantly skew the final distribution of allele or motif sizes in the truth set. After discarding variants that failed validation, the final TR truth set contained 146,640 variants, including 68,304 (47%) expansions, 68,993 (47%) contractions, and 9,343 (6%) mixed multiallelic variants.

In this study, we relied on three distinct sequencing datasets related to the CHM1 and CHM13 cell lines: 1) long read PacBio data for CHM1 and CHM13^20^, which contributed to the haploid genome assemblies used in the SynDip Benchmark and thus served as our source of truth for TR genotypes; 2) heterogeneous sequencing data from CHM13 that formed the foundation for the telomere-to-telomere (T2T) reference genome^19^; and 3) PCR-free short-read data for CHM1-CHM13 (**Methods**), which we used as the input to TR genotyping tools in order to assess their accuracy. The validation process outlined above determined the accuracy of SynDip genotypes by examining their agreement with the T2T reference genome. However, it did not incorporate the short-read data.

To find out whether the maintenance of the CHM1 and CHM13 cell lines or the process of short read sequencing itself could have distorted TR repeat counts, we examined truth set loci where ExpansionHunter, GangSTR and HipSTR agreed on the exact genotype but disagreed with the allele size in the T2T genome by 3 or more repeats. We found 96 such loci of which 51 were dinucleotide repeats, and 33 had 4bp motifs. We then manually inspected REViewer^21^ read visualizations for the subset of 19 discordant loci that had pure repeats and motif sizes ≥ 3bp. We found that for 18 out of these 19 loci, even though their SynDip allele sequences consisted of only pure repeats, the short read data clearly showed at least one interruption within their repeat sequence. Additionally, for 12 out of the 19 loci, the consensus genotype from the 3 tools was homozygous, while the SynDip genotype was heterozygous. Only one of the 19 loci fell within a SegDup region. We therefore concluded that a mixture of different causes explained these discordant loci, including read mapping and genotyping errors. Overall, with only 96 out of 146,640 variant loci being discordant, differences of more than +/- 2 repeats due to cell line drift or systematic short read sequencing errors were sufficiently rare as to be negligible.

### Truth set summary

When we applied the TR filter to all 518,281 ins/del variants within SynDip high-confidence regions, we found that 146,640 (28%) were simple tandem repeat variants that passed validation with the T2T reference (**Table 1**). 139,795 were pure and 6,845 were interrupted repeats (**Figure 2A**). Out of the 146,640 TR variants, 116,190 (79%) were mono-allelic while 30,450 (21%) were multi-allelic in the sense that the variant consisted of two alleles that differed from each other and from the hg38 reference. Overall, 60,900 out of 177,090 (34.4%) of the TR alleles in the truth set occurred at multi-allelic loci.

**Figure 2:**
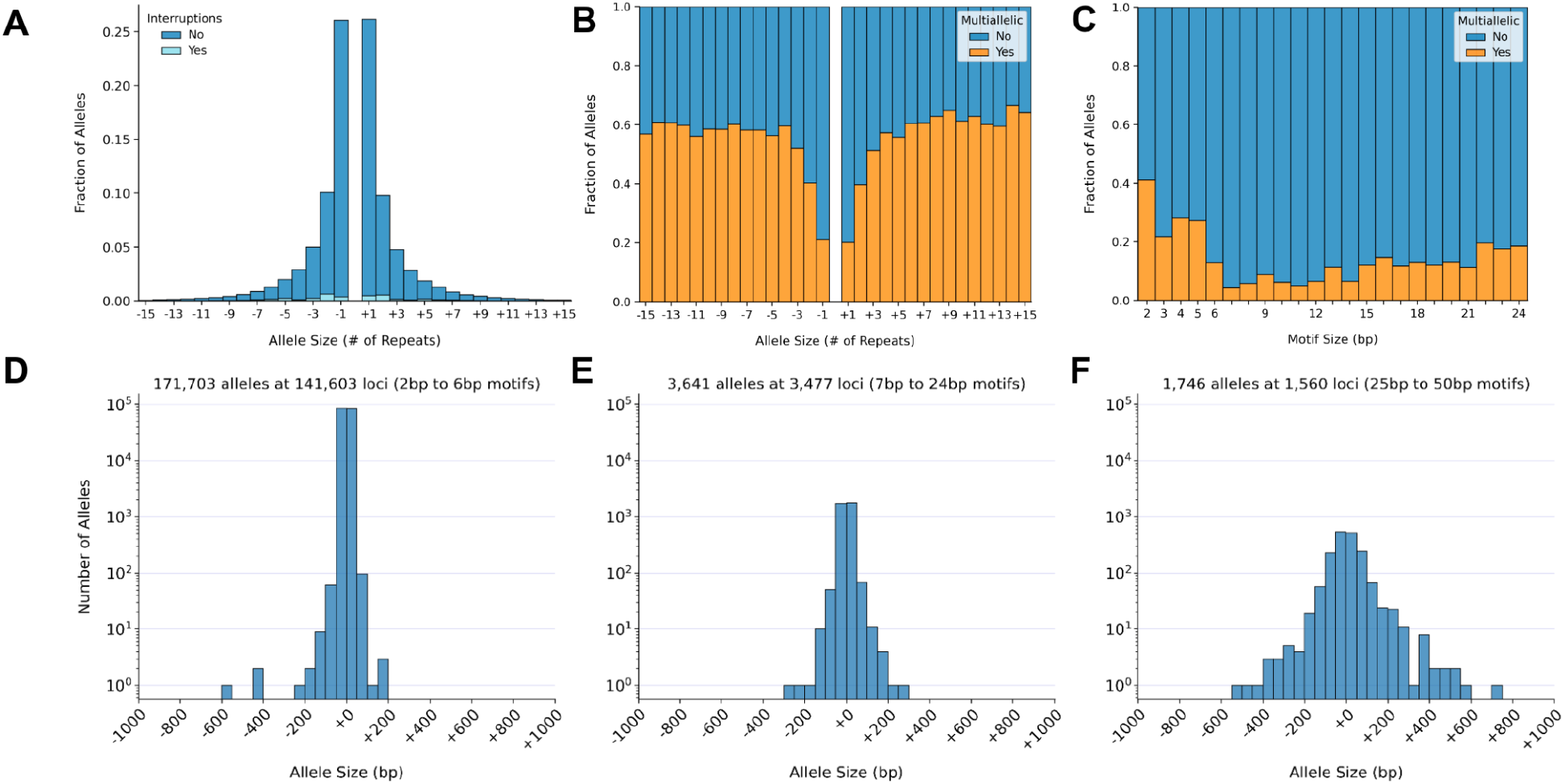
Allele size distribution. **A. Distribution of TR allele sizes.** The x axis shows truth set allele sizes minus the number of repeats in hg38 so that expansions relative to the reference are represented as positive numbers and contractions as negative numbers. The y axis shows the fraction of alleles that fall into each size bin. Darker blue represents pure repeats and light blue represents repeats with interruptions. Heterozygous reference alleles (size = +0) are not plotted. **B. Fraction of multi-allelic variants by allele size.** As in panel A, the x-axis represents true allele sizes relative to the reference. Orange represents the fraction of alleles that occur within multi-allelic loci. These are TR loci where both alleles differ from the hg38 reference and from each other. **C. Fraction of multi-allelic variants by motif size.** The x axis represents TR motif sizes ranging from 2 to 24bp. The y axis shows, for each motif size, the fraction of alleles that occurs at multi-allelic loci. This plot does not include the 1,746 alleles with motif sizes between 25-50bp. **D. Size distribution of alleles with 2-6bp motifs.** The x axis shows total allele size in base pairs, including any repeats found in the reference genome at each locus. The y axis is log-scale and shows the total number of alleles in each size bin. **E. Size distribution of alleles with 7-24bp motifs.** Same as in panel D. **F. Size distribution of alleles with 25-50bp motifs.** Same as in panel D.

In many aspects of the truth set analysis, it was important to specifically account for these multi-allelic variants and distinguish between results computed by counting alleles vs results computed by counting loci. Consequently, we were careful not to use the terms “variant” and “allele” interchangeably, but rather used “variant” to refer to a genomic site or locus as defined by its chromosome, start and end coordinates while using “allele” to refer to a specific non-reference haplotype occuring at a locus and having a unique chromosome, position, reference and alternate allele sequence.

Although in most analyses we used “allele” to refer only to non-reference haplotypes, in some analyses such as the tool evaluations, we also referred to reference haplotypes as “alleles”, explicitly noting where we did so. Finally, the size of a non-reference allele (eg. **Figure 1B**) can be measured either as an offset relative to the size of the reference allele (eg. +2 repeats or +6bp), or as its total size, which includes the reference as well as the non-reference sequence (eg. 11 repeats total). To distinguish between these two types of allele sizes, we include a + or - sign in front of the number when the size is relative, while not using a sign but adding the word “total” when specifying allele sizes that include the adjacent repeats in the reference.

In the truth set, the fraction of multi-allelic loci varied across both motif (**Figure 2C**) and allele (**Figure 2B**) sizes. Multi-allelic loci tended to have alleles that differed from the reference by more repeats (**Figure 2B**). Additionally, loci with motif sizes between 2 and 5bp were more likely to be multi-allelic than loci with other motif sizes (**Figure 2C**). The truth set expansion and contraction sizes were symmetric around the size of the reference allele (+0 on the x axis) for motifs between 2 to 6bp (**Figure 2D**), 7 to 24bp (**Figure 2E**), and 25 to 50bp (**Figure 2F**). In the tails of these distributions, 467 contractions and 523 expansions differed from the reference by more than 50bp. Conversely, 173,951 out of 177,090 (98%) of TR alleles differed from the reference by 30bp or less.

The distribution of non-reference allele sizes in the truth set was nearly identical to the distribution of repeat sizes in hg38 at truth set loci (**Figure 3A**). For loci with 2-6bp motifs, the median number of repeats was 11 in both distributions, and the median allele size was 30bp. TR loci that had fewer than 11 repeats in hg38 tended to harbor truth set expansion alleles, while reference loci with more than 11 repeats tended to harbor contractions (**Figure 3B**). Since the median number of repeats differed by motif size from a maximum of 15 repeats (for dinucleotides) to a minimum of 3 repeats (for motifs larger than 6bp), we also plotted the distributions separately for 2, 3, 4, 5, 7-24, and 25-50bp motifs in **Supplementary Figure 5A-F**. Then, to compare motif size distributions between truth set loci and TR loci in hg38 more broadly, we ran TandemRepeatFinder (TRF)^22^ on the hg38 reference and identified 1.3 million pure repeats that spanned at least 12bp, excluding homopolymers. The distribution of motifs at these hg38 loci was similar to the distribution of motifs in the truth set (**Figures 3C, D**). The proportions of some motifs such as AT and AC were noticeably larger among truth set alleles than among the hg38 loci, reflecting their relatively high mutation rates^23^. We explore TRF catalogs based on hg38 as well as mutation rates in more detail in later sections and in **Figure 7D, E**.

**Figure 3:**
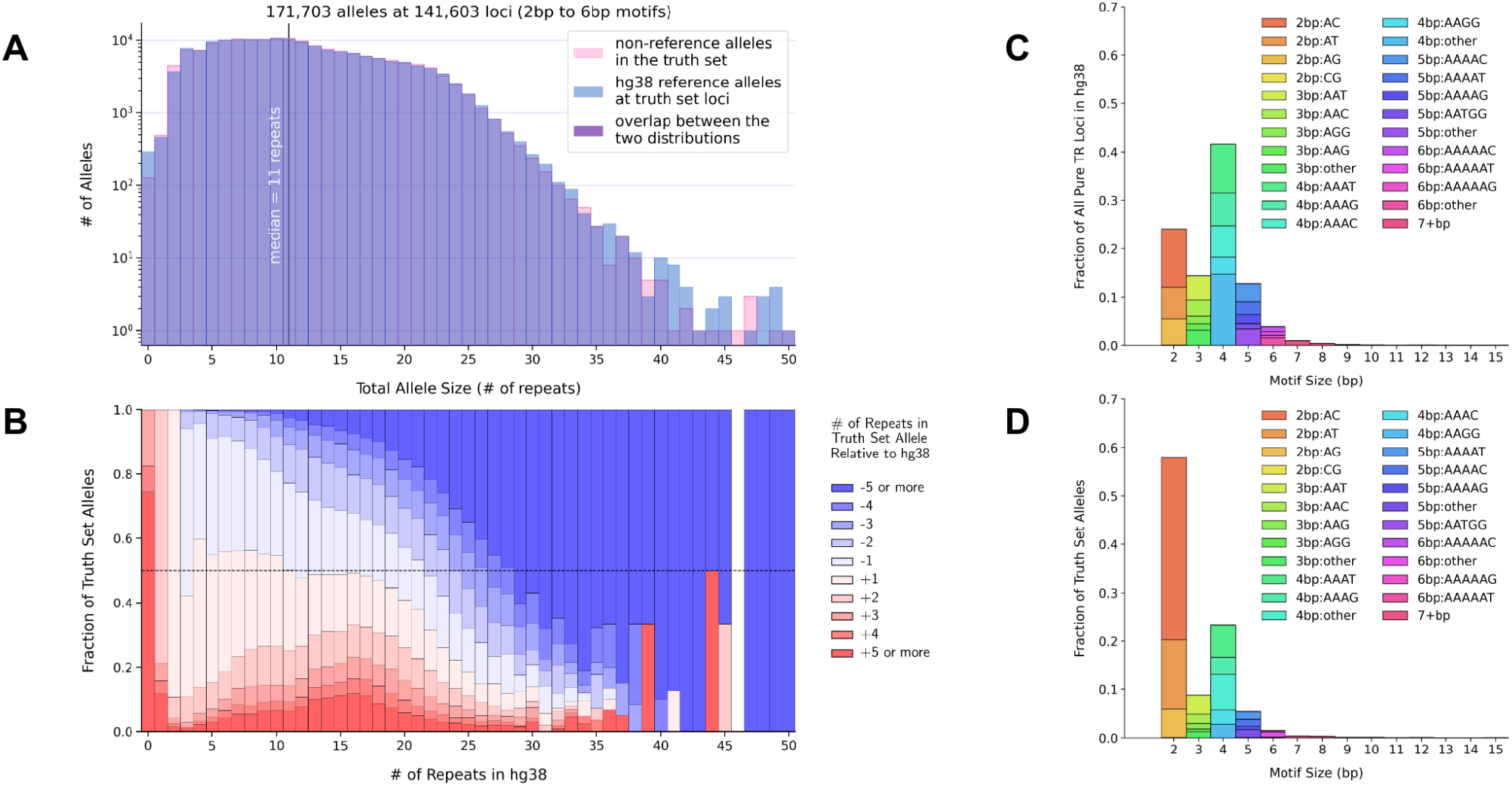
TR allele and motif sizes compared to repeat sequences in hg38. **A. Truth set allele sizes vs reference allele sizes at truth set loci with 2-6bp motifs.** The x axis shows allele sizes in terms of numbers of repeats. The y axis counts how many alleles of the given size (x axis) were found among truth set non-reference alleles (pink) or among hg38 reference alleles at truth set loci (blue). The two distributions are plotted on top of each other using semi-transparent colors so that their overlap appears purple. The vertical line at x = 11 indicates the median number of repeats, which has the same value for both distributions. For the blue distribution, the reference allele size is counted twice at multi-allelic loci so that the two distributions have the same number of counts. **B. Reference repeat sizes with 2-6bp motifs vs size of expansion or contraction at that locus.** This plot shows, for each reference locus size bin, what fraction of truth set alleles were expansions or contractions relative to the reference. The x axis is the same as in panel A. The y axis represents the fraction of the total number of alleles in each size bin. Colors show truth set allele sizes relative to the number of repeats in the reference. Darker red represents larger expansions while darker blue represents larger contractions. The horizontal dashed black line indicates where y = 0.5. **C. Motif size distribution in hg38.** For all pure repeat loci in the hg38 reference sequence that span 12 or more base pairs, this plot shows the frequency of different motif sizes and normalized motifs. Motif normalization involves taking the motif sequence and its reverse complement, computing all possible cyclic shifts of these sequences, and then selecting the cyclic shift that is alphabetically first, so that, for example, CAT, GAT, and TGA would all be normalized to ATC. The x axis represents motif sizes in base pairs. The y axis represents the fraction of all loci that have a given motif size. Colors distinguish the most frequently occurring normalized motifs within each motif size bin. **D. Motif size distribution in the truth set.** Motif sizes of all TR alleles in the truth set, shown using the same axes and colors as panel C.

### Widely-used repeat catalogs are missing substantial fractions of truth set loci

ExpansionHunter, GangSTR, HipSTR and most other tools require users to provide the exact coordinates and motifs of all TR loci to genotype. Defining this input catalog of TR loci represents a key step in any genome-wide analysis since variants occurring outside of these loci are guaranteed to be missing from the callset. Therefore, to evaluate the completeness of publicly available TR catalogs, we tested how well they captured variant loci in the truth set. Specifically, we tested three catalogs - the HipSTR catalog of 1.64 million loci, the Illumina catalog of 174,300 loci, and the GangSTR v17 catalog of 1.34 million loci. To estimate how many truth set variants would be missed by each of these catalogs, we used bedtools^24^ to compute the fraction of truth set loci that had zero overlap with the loci in a given catalog (**Methods**). We found that the HipSTR, Illumina, and GangSTR v17 catalogs respectively missed 17%, 39% and 43% of the truth set (**Table 3**).

**Table 3:** Overlap between truth set TR variants and widely-used repeat catalogs. Three existing publicly-available TR catalogs, as well as 4 new catalogs of pure repeats created as part of this truth set analysis, are compared in terms of their catalog size, motif composition, and the total number and percentage of truth set loci missed by each catalog.

| Catalog name | Catalog size:<br># of loci | Catalog size:<br>% homopolymers | Motif sizes | How it was created | # of truth set<br>loci missed | % of truth set<br>loci missed |
| --- | --- | --- | --- | --- | --- | --- |
| GangSTR catalog v17 | 1,340,266 | 36% | 1-20bp | Running TandemRepeatFinder (TRF) on hg38 with mismatch penalty = 5, indel penalty = 17 | 61,169 | 41.7% |
| Illumina catalog | 174,300 | 0% | 2-24bp | Polymorphic STR loci based on 2,504 genomes from 1kGP | 54,704 | 37.3% |
| HipSTR catalog | 1,638,945 | 51% | 1-24bp | Running TRF on hg38 using default params: mismatch penalty = 7, indel penalty = 7 | 24,254 | 16.5% |
| New catalog<br>(loci ≥ 6bp) | 4,880,610 | 0% | 2-50bp | Running TRF on hg38 using very large indel & mismatch penalties to find all pure repeats that **span at least 6bp** | 6,270 | 4.3% |
| New catalog<br>(loci ≥ 9bp) | 2,805,842 | 0% | 2-50bp | Running TRF on hg38 using very large indel & mismatch penalties to find all pure repeats that **span at least 9bp** | 7,825 | 5.3% |
| New catalog<br>(loci ≥ 12bp) | 1,343,313 | 0% | 2-50bp | Running TRF on hg38 using very large indel & mismatch penalties to find all pure repeats that **span at least 12bp** | 12,586 | 8.6% |
| New catalog<br>(loci ≥ 15bp) | 702,486 | 0% | 2-50bp | Running TRF on hg38 using very large indel & mismatch penalties to find all pure repeats that **span at least 15bp** | 24,099 | 16.4% |

We then generated new catalogs that, for a given catalog size, captured larger fractions of truth set loci even though they were not based on the truth set. To do this, we ran TRF on hg38 with very large (=1000000) mismatch and indel penalties so that it would report only pure repeats. Then, we discarded loci that spanned fewer than a minimum number of bases. By varying this minimum, we produced catalogs with 4.9 million loci spanning at least 6bp in hg38, 2.8 million loci spanning at least 9bp, 1.3 million loci spanning at least 12bp, 0.7 million loci spanning a minimum of 15bp, and so on. The catalog with 2.8 million loci missed 5.3% of the truth set while the 0.7 million catalog missed 16.4% (**Table 3**; **Supp. Figure 2**).

Since the pipeline used to create the truth set was capable of identifying TR variants in SynDip even when they had 0 matching repeats in the hg38 reference, the truth set allowed us to test the limits of a catalog-based approach to genotyping TRs. Specifically, we checked what fraction of TR variants occurred at loci with 0 matching repeats in the reference genome, and found that there were only 689 out of 157,601 (0.4%) such loci prior to validation, and 322 out of 146,640 (0.2%) that passed validation, indicating that, theoretically, a catalog based on hg38 could capture up to 99.6% of TR variants in females of European ancestry such as the SynDip sample. An additional 707 out of 146,640 (0.5%) of loci had only 1 matching repeat in the reference, and 5,082 out of 146,640 (3.5%) had 2 matching repeats. This implies that, to create the most comprehensive genome-wide TR catalog, we would need to merge a TandemRepeatFinder-based catalog generated from the reference genome with catalogs created by identifying polymorphic loci without regard to their locus size in the reference - such as the Illumina catalog of 174k loci, or loci from this truth set. This combination could capture both reference loci that are rarely polymorphic, as well as polymorphic loci which have too few repeats in the hg38 reference to be represented in TRF-based catalogs.

### Evaluating ExpansionHunter, GangSTR and HipSTR accuracy

We applied the truth set to test the accuracy of TR genotyping tools using the following steps. First, we took the 146,318 truth set loci that had more than zero repeats in the reference and converted their motifs and start/end coordinates into the repeat catalog input formats required by each tool. Next, we used these repeat catalogs to run each tool on publicly-available short read genome sequencing data from the SynDip CHM1-CHM13 sample. We then compared each tool’s output genotypes with the true allele sizes defined in the truth set by computing two per-allele accuracy metrics: strict accuracy and detailed accuracy. As illustrated in **Figure 4A**, for each locus, we compared the tool’s estimated short allele size with the true short allele size, and the tool’s estimated long allele size with the true long allele size. We counted alleles as strictly accurate when a tool’s estimate exactly matched the true allele size. Then, for detailed accuracy, we computed the difference between the estimated and true allele sizes. Unlike previous analyses where we counted only non-reference alleles, we included the reference allele in the tool comparisons when evaluating heterozygous genotypes. As a result, the tool comparisons were based on 292,636 alleles.

**Figure 4:**
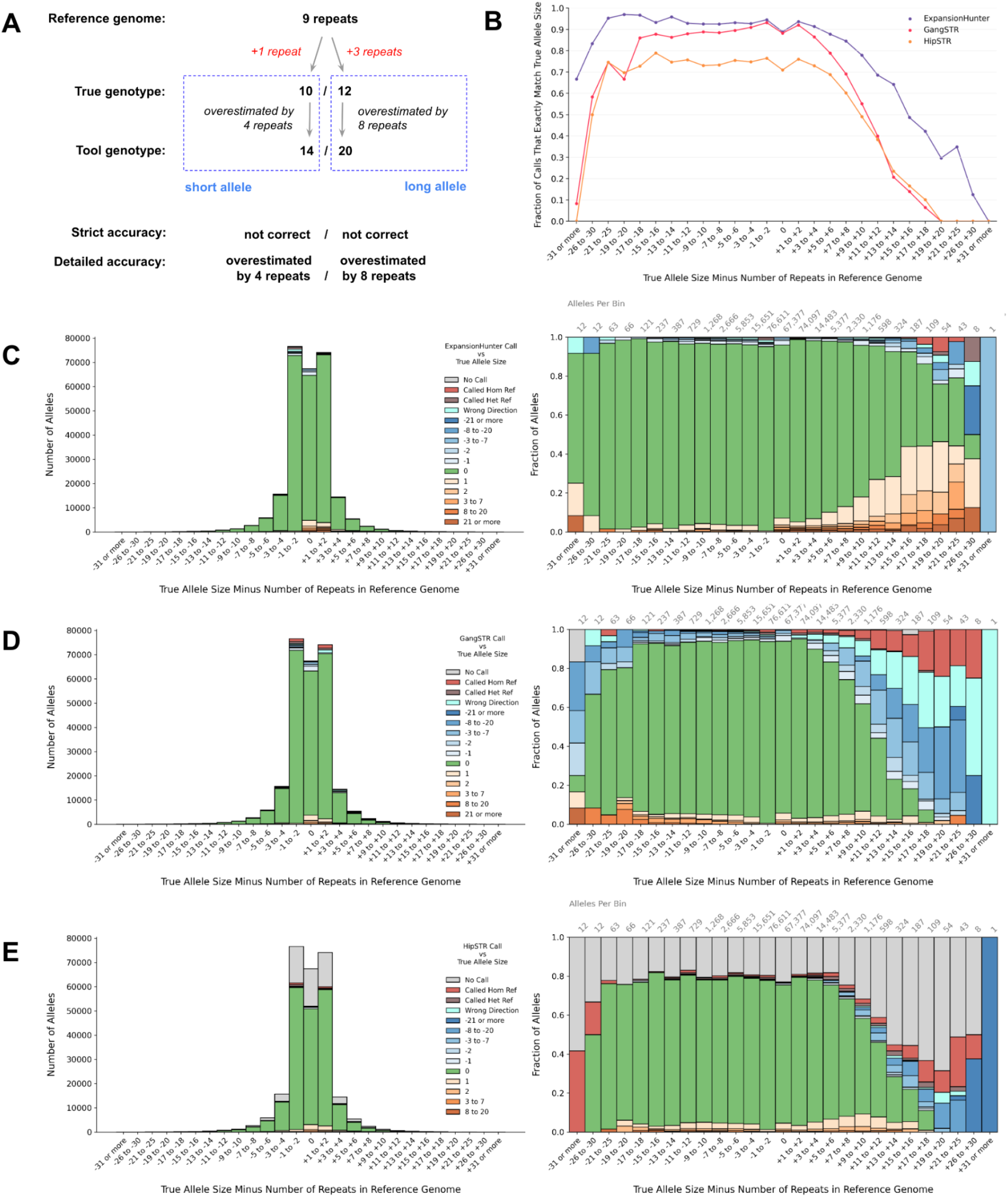
ExpansionHunter, GangSTR, and HipSTR accuracy for pure repeats with 2-6bp motifs at 30x read depth. **A. Schematic for quantifying a tool’s accuracy at a single TR locus.** When evaluating a tool’s genotype at a particular locus, there are 5 numbers to consider: the number of repeats in the reference, the two allele sizes in the truth set, and the two allele size estimates produced by the tool. In this example, the reference locus contains 9 repeats. The SynDip TR variant at the locus is multi-allelic, with the shorter allele being an expansion by +1 repeat, while the longer allele is an expansion by +3 repeats. The true genotype is therefore 10 / 12 repeats. A hypothetical TR genotyping tool estimates the genotype as 14 / 20 repeats. To quantify the accuracy of this call, we compare the tool’s long allele size with the true long allele size, and the tool’s short allele size with the true short allele size. We then define strict accuracy as whether the tool exactly matched the true allele size, and detailed accuracy as the integer difference between the tool’s estimated allele size and the true allele size. Panel B compares tools based on strict accuracy. Panels C-E use color to show detailed accuracy (for this example, the colors would be based on the numbers in the dashed blue box). In all panels, the x axis defines bins based on the difference between the true allele size and the number of repeats in the reference (labeled in red). **B. Strict accuracy by allele size.** The x axis represents true allele sizes relative to the reference. The y axis represents the fraction of alleles that each tool got exactly right. **C. ExpansionHunter detailed accuracy by allele size.** The x axis represents true allele sizes relative to the reference. The color scale represents detailed accuracy. Green indicates that the tool exactly matched the true allele size and represents the same fraction of calls as those shown in panel B. Orange represents alleles that the tool overestimated, with darker orange indicating larger errors. Blue represents underestimated alleles with darker blue indicating larger underestimates. Teal indicates that the tool called a contraction when the true allele was an expansion, or vice versa. Brown means the tool incorrectly called an allele as heterozygous reference. Red indicates when the tool called a variant locus as homozygous reference and so entirely missed the variant. Finally, gray represents loci where the tool didn’t produce a genotype (i.e. due to errors or insufficient coverage). The left-side plot shows total allele counts on the y axis while the right-side plot shows the same information but normalizing each bin to 100% and showing the fraction of alleles on the y axis. The total number of loci represented within each bar is also labeled above the right-side plot. **D. GangSTR detailed accuracy by allele size.** Axes and colors defined as in panel C. **E. HipSTR detailed accuracy by allele size.** Axes and colors defined as in panel C.

Plotting the fraction of strictly accurate alleles across different allele size bins showed that ExpansionHunter had higher accuracy than GangSTR and HipSTR (**Figure 4B**). For all three tools, accuracy declined substantially for larger allele sizes. Also, HipSTR’s accuracy was notably lower than other tools’ as it did not produce a genotype for 30,260 out of 141,603 (21.4%) of truth set loci with 2-6bp motifs, and instead produced an error message saying that the reference context upstream or downstream of the locus was too repetitive to genotype. If we compared the tools only at the 111,291 out of 141,603 (78.6%) of loci where all 3 tools did produce a genotype, HipSTR’s accuracy was similar to GangSTR’s and lower than ExpansionHunter’s (**Supp. Figure 3**). To see how ExapnsionHunter and GangSTR accuracy varied with read depth, we downsampled the SynDip read data to 20x and 10x coverage and found that ExpansionHunter’s accuracy was relatively more resilient across coverage levels (**Supp. Figure 4A**).

Next, we plotted detailed accuracy to gauge how each tools’ allele sizes differed from true allele sizes. We found that when its reported genotype was not exactly accurate, ExpansionHunter tended to overestimate the true allele size (**Figure 4C**), while GangSTR tended to underestimate it (**Figure 4D**). For HipSTR, the error distribution was dominated by missing genotypes (**Figure 4E**). We created an online plot viewer (**URLs**) to allow further exploration of detailed accuracy for these tools across different allele types and motif sizes.

### Using quality scores to post-filter TR genotypes

Short read SNV and indel calling pipelines have widely-accepted call filtering strategies such as variant quality score recalibration (VQSR)^25^. There is not yet a similar consensus on the best post-filtering approach for TR variant calls, with recent analyses relying on differing metrics and thresholds^4, 11^. GangSTR and HipSTR both output a numeric quality score (Q score) for each variant which indicates their confidence in the reported genotype. ExpansionHunter does not provide a single quality score, so we computed two different scores based on its output (**Methods**). Then, we evaluated the effect of different minimum quality score thresholds on the accuracy and false negative rate of each tool. Initially we compared tool performance on truth set loci with pure repeats and 2-6bp motifs, including loci where one or more tools did not produce a genotype and counting missing genotypes as incorrect calls (**Figure 5A**). Then, we excluded loci with missing genotypes and repeated the analysis (**Figure 5B**). For this subset of truth set loci, the best trade-off between better accuracy and higher false negative rates occurred near a minimum quality score threshold of 0.1 to 0.2 for all tools. Performance for other allele and motif sizes can be further explored using the online plot viewer (**URLs**).

**Figure 5:**
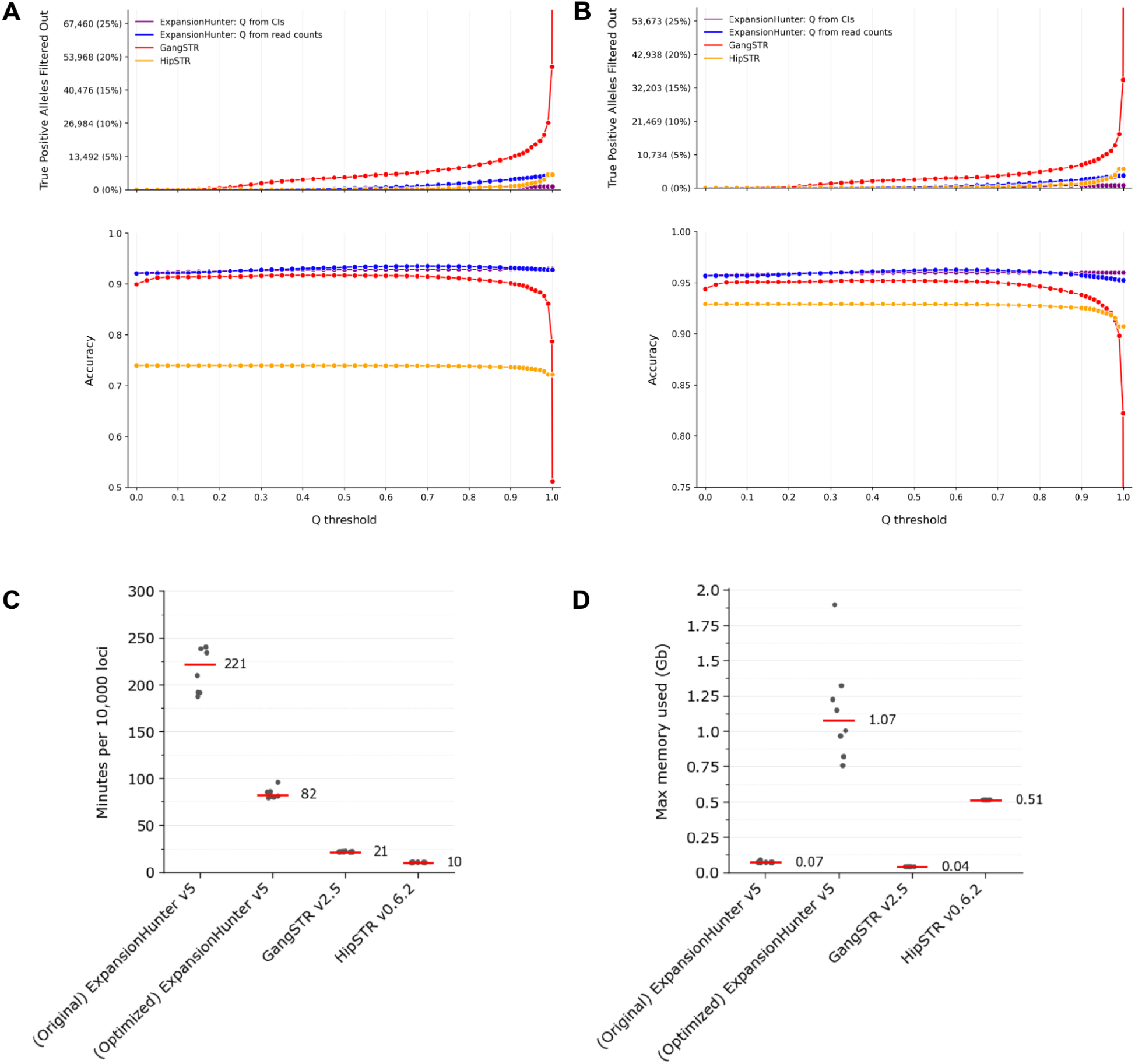
Genotype quality filters and tool performance for pure repeats with 2-6bp motifs at 30x read depth. **A. Tool performance after post-filtering using different quality score thresholds.** This plot shows how different minimum quality score thresholds affect post-filtering accuracy for calls on truth set loci with pure repeats of 2-6bp motifs and short read data with 30x coverage. The top panel shows the false negative rate, calculated as the total number or fraction (y-axis) of strictly accurate alleles that didn’t pass a given minimum quality score threshold (x-axis). The bottom panel shows the accuracy (y-axis) of alleles at each quality score threshold (x-axis). Here, accuracy is defined as (true positive alleles + true negative alleles)/(total alleles) and computed as follows: True positive alleles: the tool’s reported allele size exactly equals the true allele size and has a quality score above the minimum threshold True negative alleles: the tool’s reported allele size doesn’t equal the true allele size and has a quality score below the minimum threshold Total alleles: the total number of alleles, including those above and below the quality score threshold For GangSTR and HipSTR, the plot uses quality scores provided by the tools. Since ExpansionHunter doesn’t output a single quality score, the plot shows two different quality scores computed based on ExpansionHunter output: Q from CIs = 1/exp(4 * (confidence interval size)/(called allele size)) as in EnsembleTR^23^ Q from reads = the fraction of reads that support the allele size vs. all allele sizes considered by ExpansionHunter at the given locus. This is computed from fields in ExpansionHunter’s json output for each locus. **B. Tool performance excluding no-call loci.** This plot shows the same metrics as panel A after excluding loci where one or more tools did not produce a genotype. **C. Tool runtime per 10,000 loci.** The x axis indicates the two versions of ExpansionHunter being compared - the original Illumina version 5 and our optimized version - as well as the versions of GangSTR and HipSTR. The y axis measures runtime in minutes per 10,000 loci. For each tool, runtimes are computed for 8 different batches of truth set variant loci, with each dot representing one batch. The red lines and numerical labels indicate the median runtime across batches. **D. Memory usage.**The y axis shows peak memory usage in gigabytes for each tool. Dots represent the same 8 batches as used in panel C. Memory usage for ExpansionHunter (both the original and the optimized versions) is based on processing 500 loci per run, and may be higher for larger batches. Red lines and numerical labels indicate the median across batches.

### Comparing runtime, memory usage, and cost across TR genotyping tools

When genotyping catalogs with hundreds of thousands or millions of loci, the total required compute time can become an important factor in tool selection, particularly in cloud environments where the runtime directly affects costs. For ExpansionHunter, as for other tools, a significant fraction of overall runtime is spent reading data from disk. We therefore reasoned that adding in-memory read caching could improve performance. To test this, we modified ExpansionHunter to introduce an in-memory cache at the point where the tool fetched mismapped mate pairs from disk while processing a single locus. We also added an option to cache mismapped mates across loci on the assumption that different TR variants with the same motif would require mismapped mates to be retrieved from the same genomic intervals.

We then compared tools by runtime per 10,000 truth set loci (**Figure 5C**) and found that, when using the new --cache-mates option to enable cross-locus caching, the modified version of ExpansionHunter ran 2 to 3x faster than the original, but was still 4x slower than GangSTR and 8x slower than HipSTR. The runtime for all tools increased proportionally to the input sample’s average read depth (**Supp. Fig. 4B**). Additionally, we found that memory requirements were negligible for the original version of ExpansionHunter, as well GangSTR and HipSTR, but increased to as high as 2Gb for our modified version of ExpansionHunter when processing batches of 500 loci at a time. With --cache-mates enabled, the memory requirements would likely grow further if more than 500 loci were processed per run.

To test how these differences in runtime and memory usage affected costs per-sample in a cloud environment, we ran each tool on SynDip data with 30x coverage in order to genotype our new catalog of 2.8 million TR loci. These jobs were processed on Google’s preemptible virtual machines using a compute cluster managed by Hail Batch^26^ at an average cost of $0.022 per CPU per hour. In order to enhance caching performance of ExpansionHunter with the --cache-mates option, it was important to sort the catalog by normalized motif before dividing it into chunks of 500 loci. Additionally, we optimized the machine sizes and total numbers of loci processed per machine for each tool. We found that the lowest costs we could achieve per sample for all 2.8 million loci were $6.46 for the optimized version of ExpansionHunter, $1.15 for GangSTR, and $0.50 for HipSTR (**Methods**). We did not run the original version of ExpansionHunter v5 on the full catalog since, extrapolating the $2.66 cost of running it on 146,640 truth set loci, it would cost approximately $50/sample for the full 2.8 million loci.

### ExpansionHunterDenovo sensitivity and specificity

ExpansionHunterDenovo (EHdn) differs from other TR genotyping tools in several important ways that preclude a direct comparison. First, EHdn does not require users to specify the set of loci to genotype. Instead, the initial “profile” stage scans the read data to identify all reads whose sequence consists entirely of TR repeats. When these fully-repetitive reads have well-aligned mates, EHdn counts them as evidence for the presence of a larger-than-read-length expansion near the well-aligned mate’s location. The “profile” stage outputs a list of genomic intervals where, for each interval, it includes the motif it detected, as well as the number of fully-repetitive reads that support the presence of a repeat expansion near that location. These counts can then be used for case-control or outlier analyses in downstream stages.

Since the sensitivity and specificity of the initial “profile” stage can directly impact the false-negative and false-positive rates of the downstream tests, we evaluated the “profile” stage’s sensitivity by checking its ability to detect truth set expansions (**Figure 6A-C**). Additionally, we tested its specificity by measuring what fraction of its output genomic intervals had matching nearby expansions in the truth set (**Figure 6D**). We considered a truth set expansion as matching an EHdn output interval if they shared the same normalized motif and were no further than 600bp apart - a distance that significantly exceeded the median fragment length (327bp) of SynDip short read data.

**Figure 6:**
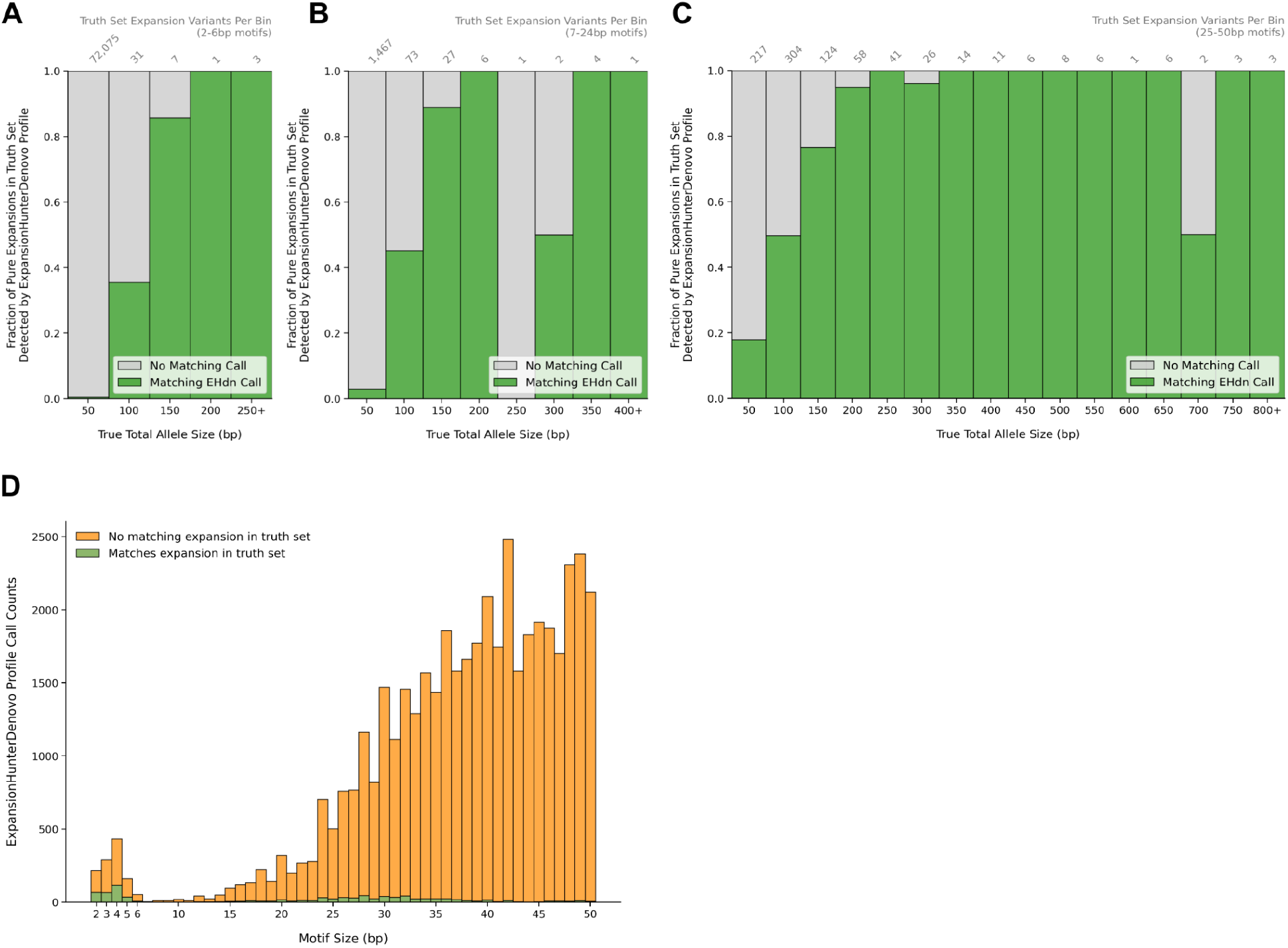
ExpansionHunterDenovo accuracy. **A. Truth set variants detected by the EHdn profile stage: 2-6bp motifs.** This plot estimates the sensitivity of ExpansionHunterDenovo’s initial “profile” stage by showing what fraction of pure repeat expansions from the truth set have a matching interval reported by EHdn. Here, an EHdn interval is considered as matching a truth set expansion if they are within 600bp of each other and have the same normalized motif. Panel A only includes truth set variants with motif sizes between 2-6bp. The x axis shows the total allele size of the truth set allele, counting the repeats in the reference as well as the variant sequence. For multi-allelic expansions, the x axis represents the size of the longer expansion since ExpansionHunterDenovo is designed to detect the largest alleles. The y axis shows the fraction of truth set variants within each bin. The number of variants represented within each bar is labeled above the plot. As expected, ExpansionHunterDenovo is most sensitive to repeats longer than the read length (150bp). The EHdn calls in this plot were generated using SynDip short read data downsampled to 30x coverage. **B. Truth set variants detected by the EHdn profile stage: 7-24bp motifs.** Same as Panel A, but only including truth set variants with motif sizes 7-24bp. **C. Truth set variants detected by the EHdn profile stage: 25-50bp motifs.** Same as Panel A, but only including truth set variants with motif sizes 25-50bp. **D. Specificity of the EHdn profile stage.** This plot estimates the specificity of ExpansionHunterDenovo’s “profile” stage by showing how many of its output intervals have a matching expansion variant in the truth set (green) including both pure and interrupted repeats, or no matching variants in the truth set (orange). As in panel A, an EHdn interval is considered as matching a truth set expansion if they are within 600bp of each other and have the same normalized motif. The x axis represents motif sizes. It shows that this initial stage of ExpansionHunterDenovo outputs many repeats that can be considered false positives, especially for larger motif sizes, leaving it to the downstream outlier detection stages to discriminate between true and false positive calls.

At 30x read depth, the EHdn “profile” stage successfully detected 321 out of 361 (88.9%) of pure truth set variants with a long allele size ≥ read length (150bp). However, 44,158 out of 45,071 (98.0%) of the genomic intervals within its output had no matching expansions in the truth set - including 860 out of 1,156 (74.4%) of intervals with 2-6bp motifs, 2,529 out of 2,662 (95%) of intervals with 7-24bp motifs, and 40,769 out of 41,253 (99%) of intervals with 25-50bp motifs. These results suggest that the EHdn profile stage has high sensitivity but poor specificity for truth set loci ≥ 150bp.

### Tandem repeats, genes, and mutation rates

We used the truth set to explore several basic properties of simple TR variation. First, we intersected truth set variants with MANE v1^27^ gene models and found that 143,953 out of 146,640 (98.2%) fell within intergenic or intronic regions (**Figure 7A**). Then, by zooming in on exonic TRs, we saw that, out of the 173 pure TR variants within coding regions, 172 (99.4%) had motif sizes that were multiples of 3bp, thereby retaining an open reading frame. This pattern continued up to motif sizes of at least 30bp (**Figure 7B**).

**Figure 7:**
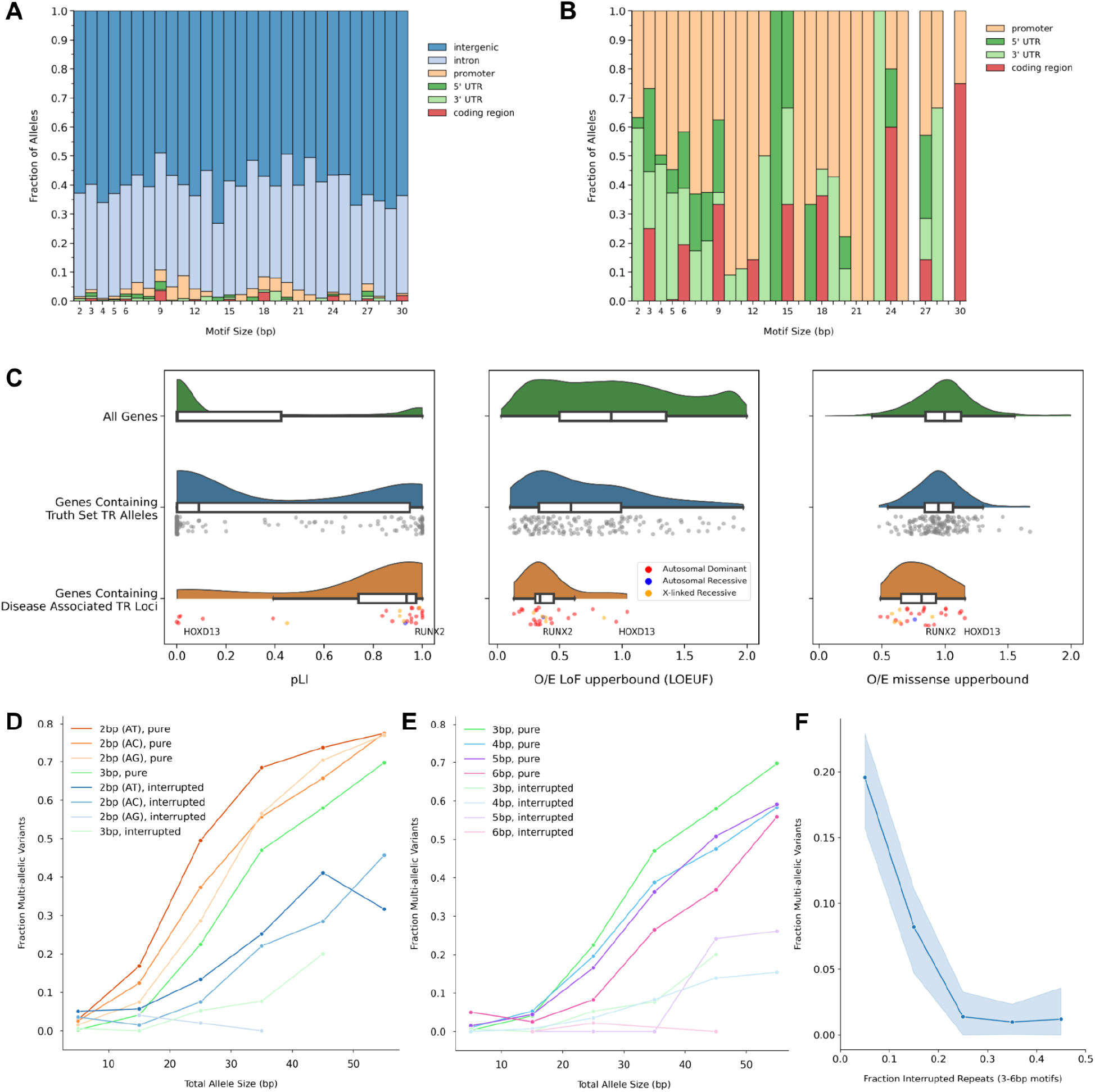
Genome-wide properties of tandem repeat variation. **A. Tandem repeats in annotated genomic regions**. This plot shows where the truth set’s pure TR variants occur relative to gene regions annotated in MANE v1. The x axis shows TR motif sizes in base pairs. The y axis shows the fraction of alleles in each bin. Colors represent the different annotated regions. **B. Tandem repeats in coding regions, UTRs, promoters.** Shows the same information as panel A, but excludes TR loci in intronics and intergenic regions in order to more clearly see the fractions for the remaining regions. **C. Gene constraint metrics vs genes that contain coding TR variants.** This plot shows 3 different measures of gene constraint on the x axis: pLI and LOEUF, and missense constraint. Above the x axis, it displays the constraint score distributions for three different sets of genes: all 19,704 genes with known constraint scores in green, 25 genes that contain known monogenic STR loci within their coding regions in orange, and 161 genes that contain pure TR variants from the truth set within their coding regions. Each dot represents a gene. In the middle track, the dot colors represent autosomal dominant (red), autosomal recessive (blue), and X-linked recessive (orange) inheritance modes of pathogenic STR expansions within those genes. Labels indicate the locations of the *HOXD13* and *RUNX2* genes which contain the two disease-associated STR loci estimated as having high constraint according to the STR-specific constraint score developed in [Gymrek 2017]. **D. Relative mutation rate by motif and allele size for 2bp motifs.** The x axis represents the true total allele size in base pairs. Orange colors represent pure repeats of different 2bp motifs (normalized so that, for example, AC, CA, GT, TG are all normalized to AC), while blue colors represent interrupted repeats. Also, repeats with 3bp motifs are shown in green for comparison (bright green for pure repeats, and light green for interrupted). The y axis represents the relative mutation rate of each bin as measured by the fraction of alleles that occurred at multi-allelic loci - ie. those where both alleles differ from hg38 and from each other. Each point in the plot represents a bin with no fewer than 20 alleles. Interrupted alleles were included in this plot if no more than 25% of the repeats within the allele sequence had interruptions (differed from other repeats in the sequence at exactly 1 position within the motif). **E. Relative mutation rate by motif and allele size for 3-6bp motifs.** Same as in panel D except that colors here represent motif sizes between 3-6bp. The brighter version of each color indicates pure repeats while the lighter color indicates interrupted repeats of that motif size. **F. Relative mutation rate by fraction of interrupted repeats.** The x axis shows the fraction of repeats within a given allele sequence that differ from the most common motif found in that sequence. For example, for an allele sequence consisting of 4 repeats CAG CAG CAT CAG, this fraction would be 0.25 due to the 1 repeat having a CAT motif. Alleles are only included when repeats differ from each other at exactly one position within the motif - such as in the 3rd position in this example. The y axis shows the relative mutation rate of each bin as measured by the fraction of alleles that occur at multi-allelic loci, similar to panel D. Each point in the plot represents a bin with no fewer than 20 alleles. The plot is based on alleles with 3-6bp motifs. The shaded region represents boot-strapped 95% confidence intervals based on variability within each point across 3, 4, 5, and 6bp motif sizes.

Next, we checked whether TR variants were enriched in genes with low constraint based on pLI^28^, LOEUF^29^, or missense constraint metrics^28^ (**Figure 7C**). We found that, out of 159 MANE v1 genes that had known pLI scores and contained truth set TR variants with pure repeats within their coding regions, 45 (28%) had high constraint based on pLI (score > 0.9) and 44 had high constraint according to LOEUF (score < 0.35). For comparison, out of 19,704 genes 3,063 (16%) were highly-constrained based on pLI and 2,971 were highly-constrained based on LOEUF (score < 0.35), indicating that SynDip TR variants were 1.8x more likely than chance to occur within genes with high loss-of-function constraint scores. This is most likely a consequence of the average coding sequence length being 1.7x longer in high-constraint (pLI > 0.9) genes than in genes with pLI < 0.9. At the same time, out of 28 known disease-associated TRs within coding regions, 18 had high pLI constraint (4.1x higher than chance), and 15 had high LOEUF constraint (3.6x higher than chance). This suggested that, while disease-associated TRs were more likely to be found within genes that had high loss-of-function constraint (particularly based on pLI), LoF-constrained genes overall were *not* depleted of in-frame TR variants in CHM1-CHM13.

Finally, we compared mutation rates of different groups of TRs by using the fraction of multi-allelic loci within each group as a proxy for that group’s relative mutation rate. This approach rested on the fact that a multi-allelic variant represented evidence for the existence of at least 3 different alleles at a locus - the allele in hg38, as well as the 2 other alleles in SynDip - while a mono-allelic variant only offered evidence for 2 alleles. Stratifying the multi-allelic fraction by total allele size and motif size reproduced known relationships between these parameters and relative mutation rates. Additionally, by separating pure TR variants from those harboring 1 or more interruptions, we found consistently lower mutation rates for alleles with interruptions (**Figure 7D**). Then, we compared multi-allelic fractions across loci stratified by the percentage of repeats that contained an interruption. We found that the relative mutation rate decreased by 3x or more as the percentage of interrupted repeats within an allele rose from 0 to 20% (**Figure 7F**).

### TR truth set completeness

The analyses in previous sections implicitly assumed that our initial ins/del variant filtering strategy identified a near-complete, or at least representative subset of simple TR variants in the SynDip genome. To evaluate this assumption and better understand what types of TR variants might have been missed, we categorized all 556,825 ins/del alleles within SynDip high-confidence regions (**Table 4**). The largest category consisted of 253,627 small indel alleles (45.5%) that did not pass TR filter criteria and had a size of 5bp or less. Next were the 188,730 ins/del alleles (33.9%) that did pass all TR filter criteria (having a motif size of 2-50bp, ≥ 3 repeats, and spanning ≥ 9bp), with 93% of these subsequently passing T2T validation and forming the truth set. Stemming from their high mutation rates, 33% of these 188,730 ins/del alleles occurred at multi-allelic loci, highlighting that they were a different class of variation from the small indel category which was only 1.3% multi-allelic, as well as from SNVs - where only 11,666 out of 3,570,146 (0.3%) of SynDip SNVs occurred at multi-allelic loci. Then, 15,537 ins/del alleles (2.8%) were found to be pure TRs with motif size > 50bp.

**Table 4:** Categorizing all ins/del alleles within SynDip high-confidence regions. We partitioned the ins/del variants within SynDip high-confidence regions into subcategories to better understand how many TR loci may have been missed by our TR filtering method. The “fraction multi-allelic” column shows the percentage of alleles within each subcategory that occurred at multi-allelic loci. We interpreted this percentage as a measure of the mutation rate, and used it to evaluate subcategories’ biological similarity to TR loci.

| Description | # of ins/del alleles | % of ins/del alleles | Fraction multi-allelic |
| --- | --- | --- | --- |
| Total ins/del alleles within SynDip high-confidence regions | 556,825 | 100.0% | 14.8% |
| Indels of size 1-5bp that did not pass TR filter criteria | 253,627 | 45.5% | 1.3% |
| TR alleles that passed all filter criteria (pre-T2T validation) | 188,730 | 33.9% | 33.0% |
| Ins/del alleles of size ≥ 6bp that did not pass TR filter criteria | 50,335 | 9.0% | 6.9% |
| TR homopolymer alleles: poly-A or T | 30,928 | 5.6% | 4.0% |
| TR alleles with pure repeats of motif size > 50bp | 15,537 | 2.8% | 25.2% |
| TR homopolymer alleles: poly-C or G | 8,157 | 1.5% | 31.3% |
| Ins/del allele within a multi-allelic variant where only one of the alleles passed TR filter criteria | 4,930 | 0.9% | 100.0% |
| TR allele that failed filter because the variant sequence ended in a partial repeat (eg. *chr6:71189213 CAGCAGCA > C*) | 3,192 | 0.6% | 14.9% |
| TR alleles that were excluded for other reasons | 1,389 | 0.2% | 33.5% |

Although these were excluded from our analysis, 25.2% of them occurred at multi-allelic loci, corroborating their biologic similarity to truth set TRs. At the other end of the motif size scale, 30,928 ins/del alleles (5.6%) represented poly-A homopolymer repeats and 8,157 (1.5%) represented poly-C repeats that were detected by the TR filter and intentionally excluded from the analysis. The multi-allelic fraction of poly-C homopolymers was 31.3% - matching that of truth set TRs. However, only 4% of the poly-A repeat loci were multi-allelic - a surprisingly low proportion that may be due to the intentional exclusion of poly-A runs ≥ 10bp by the SynDip pipeline^17^.

The only other category that contained more than 1% of ins/del alleles was the set of 50,335 ins/del alleles (9%) that were 6bp or larger and did not pass TR filter criteria. Out of the categories listed in **Table 4**, this is the only one that could contain complex TR variants missed by our filtering method, including those with complicated interruption patterns or compound repeats. As a result, we considered the size of this category to be a conservative upper-bound on the number of complex TR alleles missed by our filter. Since the multi-allelic fraction of this category was 7%, we expected less than ¼ of these alleles to represent TRs with a similar mutation rate as the TRs in the truth set.

## Discussion

Generating a genome-wide truth set for 146,640 TR variants provided us with new insights into the strengths and weaknesses of TR genotyping tools as well as the degree to which existing genome-wide TR catalogs remain incomplete. Previous studies used haploid genome assemblies for in-depth surveys of genome-wide variation and benchmarking of tools with a focus on structural variants, SNVs, indels, mobile elements, and complex VNTR sequences^17, 30–32^. We instead focused on simple repeats and existing tools for genotyping them using short read sequencing data. Through our analysis, we found that:

- ExpansionHunter is more accurate than GangSTR and HipSTR on truth set loci. Additionally, ExpansionHunter errors are more balanced between over- and underestimating the true allele size, while GangSTR is more likely to underestimate expansions (**Figure 4C, D**). We also found that, on genome-wide catalogs, ExpansionHunter is significantly slower than GangSTR and HipSTR.
- Starting with a larger TR catalog, such as our new hg38 catalogs of 2.8 million or 1.3 million TRs (**URLs**), should allow genome-wide studies to capture at least 50% more simple TR variants per individual (regardless of tool choice) as compared to available catalogs for GangSTR and ExpansionHunter (**Supp. Figure 2**).
- The initial TR discovery stage of ExpansionHunterDenovo has 89% sensitivity for alleles larger than 150bp, but has low specificity (**Figure 6**).

Additionally, we examined constraint scores of genes that harbored known disease-associated TR loci within their coding regions and concluded that pLI scores, more so than LOEUF or missense-Z scores, could be a useful metric for prioritizing candidate TR loci when investigating potential novel disease associations, particularly while TR-specific constraint metrics^33^ remain available only for a subset of coding loci. Our measurements of relative TR mutation rates for different allele and motif sizes were broadly consistent with previous studies^23, 34, 35^. We additionally compared mutation rates of interrupted vs. pure repeats and found that, consistent with expectation^36–38^, mutation rates dropped precipitously as the number of interruptions within the repeat sequence increased (**Figure 7F**). Finally, we found that the genome-wide distributions of TR sizes were nearly identical in the CHM1, CHM13, and hg38 genome assemblies (**Figure 3A**). Since the truth set represented variant loci relative to the reference sequence, we interpreted the similarity between allele size distributions as an indicator of a general similarity of simple TR allele size distributions across individuals with similar ancestry rather than just a consequence of CHM1 having been previously used to close gaps within repetitive regions of the GRCh38 reference^39^.

Similar to its use in previous studies which focused on SNV and indel variants^17^, the Synthetic Diploid Benchmark provided us with a source of truth for TRs that was orthogonal to short read data. This truth set overcame many of the limitations that exist for other ways of evaluating tool accuracy such as PCR validation (which has low throughput), simulated data (which doesn’t capture the full complexity of real sequencing data), Mendelian violation rates (which are more informative about specificity than sensitivity since a tool that missed all variants would register zero Mendelian violations), and consensus callsets across multiple tools (which leave true allele sizes unclear). Additionally, since all the underlying data and code are publicly available without restrictions, these benchmarks can be broadly applicable to future tool comparisons and continued tool development.

That being said, our approach to generating the TR truth set had several key limitations. First, it was based on SynDip data from a single female sample of European ancestry. As a result, it did not contain chrY variants, and missed an unknown number of loci that are polymorphic among other individuals within the global human population. Even within this one sample, we excluded multiple categories of tandem repeats that, for various reasons, did not pass our filtering criteria. The largest excluded categories represented homopolymers, as well as repeats with motif sizes > 50bp. Also, we excluded variation within the SynDip low-confidence regions which covered 0.32 out of 3.03Gb (10.5%) of autosomal + chrX sequence in hg38. Had we included them, we would have identified an additional 10,227 TRs with 2-50bp motifs, of which 8,526 (83%) would have subsequently passed validation using the T2T reference.

The analysis of interrupted repeats was limited to simple interruption patterns where a single position within the motif varied across repeats. The TR filter, as currently implemented, was not capable of detecting more complicated patterns, including those seen at the *RFC1* locus^40, 41^ where a single allele could include alternating repeats of AAAAG, AAGGG, ACAGG or other such motifs.

For this initial analysis, we focused on four widely-used tools designed for short read data. Adding more STR and VNTR genotyping tools, as well as tools like GATK HaplotypeCaller^25^ and DeepVariant^42^, would be helpful for comparing their strengths and behaviors on TR loci. Although exome sequencing data is available for CHM1-CHM13, we did not include its analysis in this manuscript. However, tool accuracy plots for exome data are available in our plot viewer (**URLs**). To our knowledge, neither PacBio HiFi nor Oxford Nanopore (ONT) data is available for the CHM1-CHM13 sample, limiting our ability to evaluate TR tools designed for the most recent long-read sequencing technologies.

Generating high-quality genotypes for simple repeats in a larger and more diverse set of haploid assemblies would overcome many of the limitations of our current study. First, it would lead to a more complete catalog of polymorphic TR loci and enable further validation of TR genotyping tools for short read data. Additionally, it would provide an orthogonal truth set for measuring the accuracy of TR genotyping tools designed for HiFi PacBio and ONT data, as well as novel sequencing technologies such Ultima^43^. We look forward to more haploid assemblies and TR truth sets becoming available thanks to rapid ongoing progress in long read sequencing technologies, assembly methods, and projects such as the Human Genome Structural Variation Consortium (HGSVC) and the Human Pangenome Project^44^.

In conclusion, our study demonstrated the utility of an orthogonal genome-wide truth set for measuring the accuracy of TR genotyping tools, evaluating the completeness of TR catalogs, and exploring the overall properties of TR variation.

## Methods

### URLs

Interactive plot viewer for exploring tool accuracy across different parameters: https://broadinstitute.github.io/str-truth-set/html/tool_comparison_viewer.html

IGV.js-based^45^ genome browser for TR truth set tracks: https://broadinstitute.github.io/str-truth-set/html/tgg_viewer.html

TR catalog with 2.8 million pure repeats in hg38 that captures 95% of truth set variants: https://storage.googleapis.com/str-truth-set/hg38/ref/other/repeat_specs_GRCh38_without_mismatches.sorted.trimmed.at_least_9bp.bed.gz

TR catalog with 1.3 million pure repeats in hg38 that captures 91% of truth set variants: https://storage.googleapis.com/str-truth-set/hg38/ref/other/repeat_specs_GRCh38_without_mismatches.sorted.trimmed.at_least_12bp.bed.gz

Optimized ExpansionHunter: https://github.com/bw2/ExpansionHunter

TR filter code:

https://github.com/broadinstitute/str-analysis/blob/main/str_analysis/filter_vcf_to_STR_variants.py

Code used to generate the TR truth set and perform all downstream analyses:

https://github.com/broadinstitute/str-truth-set

Results from intermediate steps of the TR truth set creation and analysis pipeline: https://storage.googleapis.com/str-truth-set

### Data

We downloaded Synthetic Diploid Benchmark variant genotypes (full.38.vcf.gz) and high-confidence regions (full.38.bed.gz) from

https://github.com/lh3/CHM-eval/releases/tag/v0.5, then used the bedtools intersect command ^24^ with -f 1 to subset these variants to those within high-confidence regions.

Short read genome data for the CHM1-CHM13-2 sample (ERR1341796) is available from the Broad Institute:

https://console.cloud.google.com/storage/browser/broad-public-datasets/CHM1_CHM13_WGS2

It was sequenced on the Illumina HiSeq X Ten instrument using a PCR-free protocol with paired-end 151bp reads. It has 40x depth of coverage and a median DNA fragment length = 327bp (+/- 67 MAD).

Exome data for the CHM1-CHM13-2 sample is also available from the the Broad institute:

https://console.cloud.google.com/storage/browser/broad-public-datasets/CHM1_CHM13_WES

The exome was sequenced on an Illumina HiSeq X Ten instrument with paired-end 151bp reads. It has 85x mean target coverage and a median DNA fragment length = 393bp (+/- 87 MAD).

### Code

The TR filtering algorithm, which takes a single-sample VCF and outputs the subset of ins/del variants that represent TR expansions or contractions, is implemented in *filter_vcf_to_STR_variants.py* in the https://github.com/broadinstitute/str-analysis repo.

All steps for generating the TR truth set (as well as all results and figures in this paper) are encoded in *run_all_steps.sh* within the https://github.com/broadinstitute/str-truth-set repo. This script consists of 5 sub-steps, named step A through E:

Step A: can run locally on any computer with a recent version of MacOS and the necessary software installed (python3, bedtools, gatk, bgzip, tabix). It downloads SynDip, hg38 and T2T reference data, installs required python packages, and then proceeds to run the commands needed to produce the TR truth set, validate it against the T2T reference, and generate downstream statistics about its overlap with genes and widely-used TR catalogs.

Steps B through E require write access to a Google storage bucket, as well as the Hail Batch^26^ system which we used to run TR genotyping tools and generate figures on a cluster of Google cloud machines. Although these steps are not directly reproducible without paid accounts on these systems, all intermediate and final outputs are publicly available @ https://console.cloud.google.com/storage/browser/str-truth-set/

Step B: runs the *tool_comparison/scripts/convert_truth_set_to_variant_catalogs.py* script to convert the truth set into the repeat catalog input formats required by ExpansionHunter, GangSTR and HipSTR. Then, it uploads these and other files to the Google storage bucket.

Step C: launches Hail Batch pipelines to 1) downsample the SynDip short read genome data from its original 40x coverage to lower coverage levels, and 2) run ExpansionHunter, GangSTR, HipSTR, and ExpansionHunterDenovo for each coverage level.

Step D: combines results from step C into the tables used for downstream analyses. This is also the step that computes concordance between tool results and truth set TR genotypes as outlined in **Figure 4A**.

Step E: Generates all figures and tables.

### Filtering SynDip insertions and deletions to TRs

We filtered the variant calls published by the Synthetic Diploid Benchmark (full.38.vcf.gz) to those within SynDip high-confidence regions (full.38.bed.gz) by running bedtools intersect -f 1. Then we ran *filter_vcf_to_STR_variants.py* to identify insertions and deletions that represent TR expansions or contractions, using the command line:

`python3 -u -m str_analysis.filter_vcf_to_STR_variants -R`

`./ref/hg38.fa --allow-interruptions`

`only-if-pure-repeats-not-found --min-str-length 9`

`--min-str-repeats 3 --min-repeat-unit-length 2`

`--max-repeat-unit-length 50 --write-bed-file`

`--write-vcf-with-filtered-out-variants step1.high_confidence_regions.vcf.gz`

We subsequently reran this command to detect homopolymers by changing --min-repeat-unit-length 2 to --min-repeat-unit-length 1

The following details about the TR filtering algorithm did not fit into the main text. When an allele with interruptions passed TR filter criteria, the algorithm took the most common motif within the variant sequence to be the TR motif. For example, “CAG” would be the most common motif in a C > CATCAGCAG insertion since “CAT” occurs only once.

Then, the algorithm would look for additional CAG repeats in the reference, and thereby derive the reference start and end coordinates of the locus. Although the algorithm would allow repeats in the reference to have a similar interruption pattern (“CAN” in the example, where N represents any base), it would require that the first and last repeat copy in the reference exactly match the common motif (“CAG”). This rule was intended to prevent over-extending the reference locus and assumed a strict meaning of “interruption” as a temporary departure from the most common repeat motif.

For multi-allelic TR variants, these additional rules sometimes led to an edge case where each of the two alleles passed the TR filter criteria, but differed in their motifs, reference start/end coordinates, or position within the motif that had interruptions. In SynDip, the filter discarded 227 such variants. Additionally, the algorithm checked whether nearby ins/del variants that passed TR filter criteria resulted in reference TR loci that overlapped each other. Since, in these cases, it was unclear which ins/del variant represented the true TR genotype for that locus, the ins/del variants were discarded. In SynDip, this led to the exclusion of 449 TR variants.

Finally, when checking a variant sequence for dinucleotide motifs with interruptions, the filter used stricter rules than simply allowing one position within the dinucleotide motif to vary across repeats. Specifically, it required ⅘ of repeats to exactly match the most common motif. For detecting homopolymers with interruptions, similar constraints were added, requiring 9 out of 10 bases to be pure homopolymer stretches and disallowing interruptions that spanned two or more bases in a row. Additionally, when extending dinucleotide or homopolymer repeats into the reference context, the filter did so based only on pure repeats, and did not extend these motifs past interruptions.

### Validating TRs using the telomere-to-telomere reference

We validated variants that passed the TR filter by lifting them over to the T2T reference using the GATK LiftoverVcf tool and the UCSC hg38-chm13v2.chain file. Then, since one of the two haplotypes within the SynDip CHM1-CHM13 sample is also the basis of the T2T reference genome (CHM13), we checked that at least one of the two alleles in each variant matched the T2T reference sequence, as implemented in *scripts/filter_out_discordant_variants_after_liftover.py*. To avoid a dependence on phasing accuracy, we did not take SynDip phasing information into account while performing validation.

To overcome limitations in the way LiftoverVcf processed TR variants, we implemented pre- and post- processing steps. The first limitation of GATK LiftoverVcf was that it distorted TR allele sizes for the many loci where hg38 had a different number of repeats than the corresponding locus in T2T. For example, there are 6 GAA repeats at the hg38 *FXN* locus (chr9:69,037,287-69,037,304), but 9 GAA repeats at the same locus in T2T (chr9:81,210,835-81,210,861). As a result, a hypothetical chr9:69,037,286 A > AGAA insertion of +1 repeat represented a total allele size of 7 x GAA repeats based on hg38. To preserve this allele size in the T2T context would require converting this variant from an insertion of +1 x GAA to a deletion of -2 x GAA repeats. Instead, GATK LiftoverVcf naively mapped this variant to chr9:81,210,843 A>AGAA, producing a new TR allele size of 10 x GAA. To avoid these distortions, we ignored the ref. and alt. allele fields in the post-liftover VCF and instead relied only on the variants’ T2T start position when evaluating concordance, as described below.

Another limitation was that more than 70% of deletions and over 95% of mixed multi-allelic variants (consisting of 1 deletion and 1 insertion allele) failed liftover with an “IndelStraddlesMultipleIntevals” error due to the deletion spanning boundaries between liftover chain intervals. To mitigate this, after applying the TR filter and before running the hg38 ⇒ T2T liftover, we converted all mono-allelic deletions to SNVs that had the same start position as the deletion and an arbitrary base for the alternate allele (see *scripts/convert_monoallelic_deletions_to_snvs_for_liftover.py*). This allowed us to use LiftoverVcf’s hg38 ⇒ T2T coordinate mapping functionality without triggering the “IndelStraddlesMultipleIntevals” error, and since our validation procedure relied only on the repeat counts which had already been computed by the TR filter, we could restore the original reference and alternate alleles later, post-validation.

To evaluate each variant’s concordance with T2T after the hg38 ⇒ T2T liftover, we first took the repeat motif identified by the TR filter pre-liftover and computed the total number of repeats of that motif in the T2T reference immediately to the left and right of the variant’s T2T position. For this step, we reused the algorithm implemented within the TR filter where it extended repeats of a given motif to the left and right of a variant’s position. We then considered a variant as passing if the number of repeats in the T2T reference sequence was within +/-2 of the number of repeats in the variant’s short or long allele (or both) as identified by the TR filter pre-liftover. We allowed the 2-repeat margin to account for TR stutter during CHM13 cell-line maintenance and sequencing.

In the end, we used the UCSC chm13v2-hg38.chain file to lift variants back over to hg38 and ran the *scripts/check_vcf_concordance_before_vs_after_liftover.py* script to make sure that all variants still had the same hg38 position after hg38 ⇒ T2T ⇒ hg38 liftover. Variants that passed all the above validation steps were added to the truth set.

Multi-allelic deletions and mixed multi-allelic variants that failed the hg38 ⇒ T2T liftover due to the “IndelStraddlesMultiplelIntevals” error were then appended to the truth set without validation, since this error prevented their comparison to the T2T reference.

### Running TR genotyping tools and evaluating results

We ran ExpansionHunter, GangSTR and HipSTR using default parameters except for the addition of --def-stutter-model and --min-reads 5 when running HipSTR, similarly as described in recent publications^46, 47^. Also, when running ExpansionHunter on the downsampled 10x coverage sample, we added --min-locus-coverage 3. For the ExpansionHunterDenovo profile command, we used --max-unit-len 50.

The outputs of these tools were combined into tab-delimited tables by running the *str_analysis/combine_str_json_to_tsv.py* script. This script is also where we computed an allele quality score for ExpansionHunter using the same formula as EnsembleTR^23^: Q_CI = 1/exp(4 * (confidence interval size)/(total allele size)).

Additionally, we computed a second allele quality score for ExpansionHunter by calculating the number of reads that support the given allele size divided by the total number of reads ExpansionHunter considered during genotyping. These numbers are derived from ExpansionHunter’s json output fields CountsOfSpanningReads, CountsOfFlankingReads, and CountsOfInrepeatReads. Each field contains a list of 2-tuples which pair an allele size with the number of spanning, flanking and in-repeat reads that support it. We found that allele sizes within the CountsOfInrepeatReads

2-tuples plateaued when they approached read length. We therefore considered the CountsOfInrepeatReads as supporting any allele size that exceeded 80% of the read length. This calculation produces the “FractionOfReadsThatSupportsGenotype: Allele 1 (or Allele 2)” field in *str_analysis/combine_str_json_to_tsv.py* output. Later, for better empirical performance, we rescale this fraction by mapping any value > 0.15 to 1 and dividing any value between 0 and 0.15 by 0.15 to produce a Q score between 0 and 1.

In step D of *run_all_steps.sh*, we annotated the truth set table with the genotypes reported by ExpansionHunter, GangSTR and HipSTR by running the *tool_comparison/scripts/add_tool_results_columns.py* and *tool_comparison/scripts/add_concordance_columns.py* script for each tool, followed by *tool_comparison/scripts/combine_all_results_tables.py*. Tool results were joined with the truth set based on locus IDs derived from the chromosome, start, end and motif of each TR variant. Concordance was computed as described in **Fig. 4A**.

Additionally, we ran *tool_comparison/scripts/intersect_expansion_hunter_denovo_results_with_truth_set.py* with --window-size 600 to determine overlap between EHdn calls and truth set variants. This script parsed the EHdn “profile” command’s output locus.tsv file where each row included a motif, chromosome, start and end coordinates. EHdn calls were considered as matching a truth set expansion variant if they occurred within 600bp of each other and had the same motifs after normalization (taking the version of the motif that’s alphabetically first across all cyclic shifts of the motif and its reverse complement).

Finally, step E used the combined tables to generate the figures, tables, and statistics in this manuscript.

### Tool cost optimization on Hail Batch

To estimate overall costs per sample, we ran each tool on the 30x coverage SynDip sample to genotype our new catalog of 2.8 million loci. We converted this catalog to the appropriate input formats for each tool and reused the same command lines as when measuring tool accuracy - mostly using default parameters except for HipSTR where we added --min-reads 5 and --def-stutter-model.

To parallelize execution across multiple machines, we used a cluster of preemptible Google Cloud VMs managed by Hail Batch. After a fixed data localization cost per job, the costs scaled linearly with time used per CPU. The system provided machines with between 0.25 and 16 CPUs at a cost of $0.022 per CPU per hour, and included 3.75 Gb of RAM per CPU with the option to increase this to 7 Gb per CPU for a 10% increase in cost. Preemptible machines were most cost-effective for jobs that took no more than a few hours to complete as this reduced the chance of the jobs being preempted and needing to restart from the beginning. Therefore, we split the full catalog into smaller chunks that completed quickly. Additionally, we experimented with different machine sizes and within-machine parallelization in order to maximize utilization of CPUs and memory by each tool.

For HipSTR, we processed the full catalog in chunks of 10,000 loci per HipSTR instance, using 4-CPU machines with extra memory (highmem) and running 32 HipSTR processes in parallel on each machine (8 per CPU). By parallelizing across 3 such machines, the overall pipeline completed after 1 hour and cost $0.50 total. For GangSTR, we used 2-CPU machines with a standard amount of memory, and ran 16 GangSTR processes in parallel on each machine (8 per CPU). By using 18 machines, the pipeline completed in 1 hour at a cost of $1.15. We ran the optimized version of ExpansionHunter on chunks of 500 loci at a time. To increase cache hit rates, we arranged the chunks to contain loci with the same motif whenever possible. When using 16 4-CPU machines and running 12 ExpansionHunter processes in parallel per machine (3 per CPU), the pipeline took 8 hours and cost $6.46. Further optimization could have been achieved by recording processing times of individual loci via the new --record-timing option and then removing loci that took ExpansionHunter the longest to genotype. For the original Illumina version of ExpansionHunter v5, we found the cost was lowest when running 4 processes per CPU on machines with 2 CPUs and standard memory. We did not run this on the full catalog of 2.8 million loci, but extrapolated the cost from processing 146,640 truth set loci which came out to $2.66 x 2,800,000 / 146,460 ≈ $50.

### Testing completeness of widely-used repeat catalogs

We evaluated how many truth set variant loci were missing from widely-used TR catalogs for hg38, including the GangSTR v17 catalog downloaded from https://github.com/gymreklab/GangSTR, the lllumina catalog of 174k polymorphic loci downloaded from https://github.com/Illumina/RepeatCatalogs, and the HipSTR catalog from https://github.com/HipSTR-Tool/HipSTR-references.

This comparison was implemented in *scripts/compute_overlap_with_other_catalogs_using_bedtools.sh* script as part of Step A within *run_all_steps.sh* . That script used the following bedtools command to count the number of loci in the truth set that did not overlap any loci in a given catalog:

`bedtools subtract -a $truthset_bed -b $catalog_bed -A | wc -l`

### Generating new TR catalogs

To generate a comprehensive catalog of pure TR repeats in hg38, we ran TandemRepeatFinder v4.09 with Match=2, Mismatch Penalty=1000000, Indel Penalty=1000000, PM=80, PI=10, Minscore=8, MaxPeriod=2000 using the following command line:

`trf catalog.txt 2 1000000 1000000 80 10 8 2000 -ngs -d`

Then, we trimmed the end coordinate of each locus so that the number of base pairs was an exact multiple of the motif size. Finally, we filtered the results to produce a catalog of 2.8 million loci that spanned at least 9bp, a catalog of 1.3 million loci that spanned at least 12bp, and so on.

## Acknowledgements

The authors would like to thank Marina and Joseph Weisburd for all the ways in which they made this manuscript possible. Thank you to the Hail team for tools that made the analysis much easier than it would have been otherwise. Also, thank you to Hope Tanudisastro, Nehir Edibe Kurtas and Katherine Chao for helpful discussions.

Funding was provided by NIH/NHGRI grants UM1HG008900 and U01HG011755.

---

**Supplementary Figure 1:**
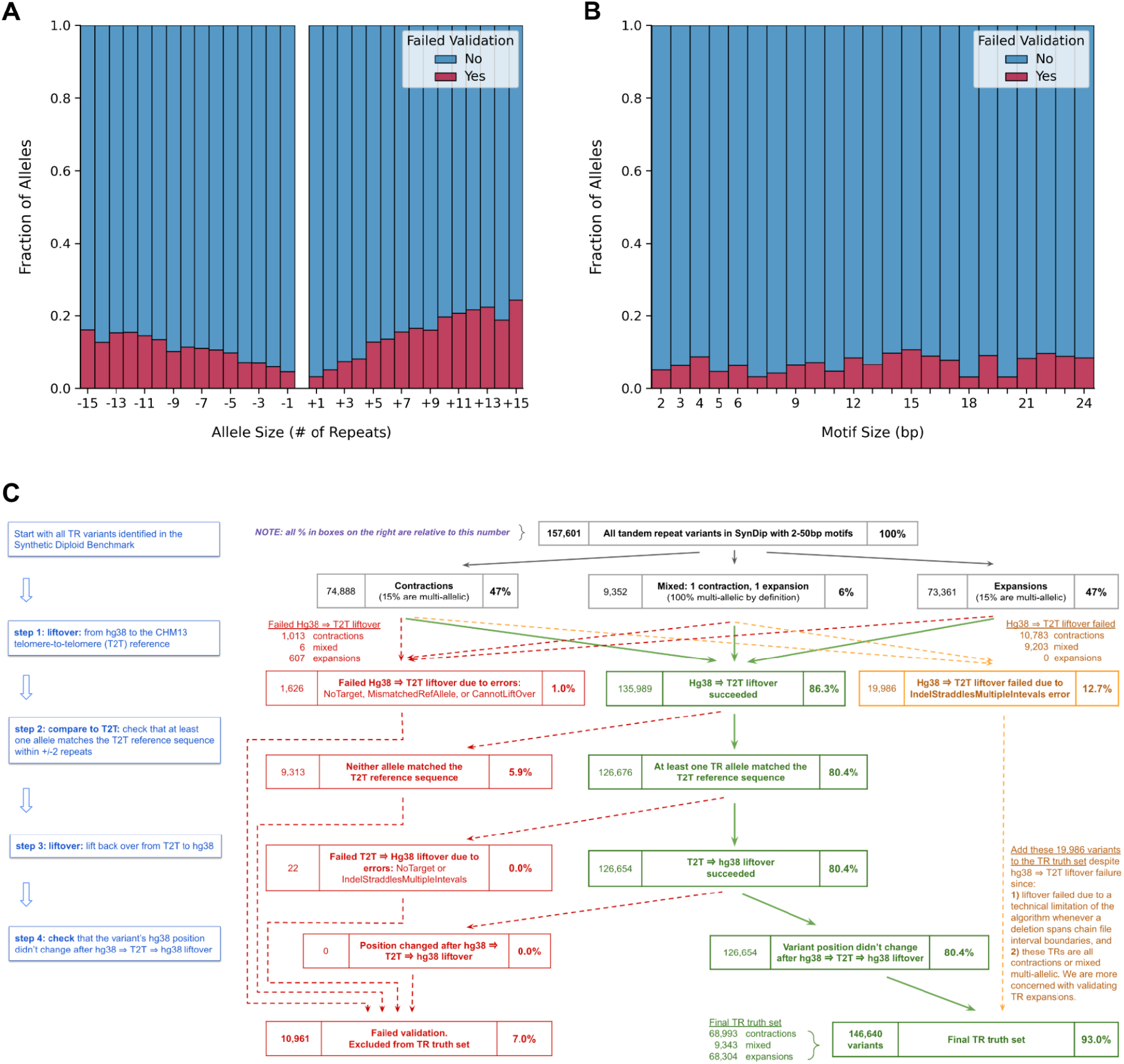
Validation of TR variants via liftover to the telomere-to-telomere (T2T) reference genome. **A. Distribution of allele sizes for loci that failed validation.** The x-axis represents SynDip allele sizes relative to the hg38 reference. Red represents the fraction (y-axis) of alleles that occurred at loci which failed validation. **B. Distribution of motif sizes for loci that failed validation.** The x axis represents TR motif sizes ranging from 2 to 24bp. The y axis shows, for each motif size, the fraction of alleles that occurred at loci which failed validation. **C. Validation procedure flowchart and statistics.** This flowchart shows the 4 steps (blue) used to validate TR truth set variants via comparison to the T2T reference sequence, as well as the number of variants that passed (green) or failed (red) each step. It also shows the contractions and mixed multi-allelic variants that could not be validated (orange) due to technical limitations of the liftover algorithm. In the end, 10,961 (7%) of TR variants failed validation.

**Supplementary Figure 2:**
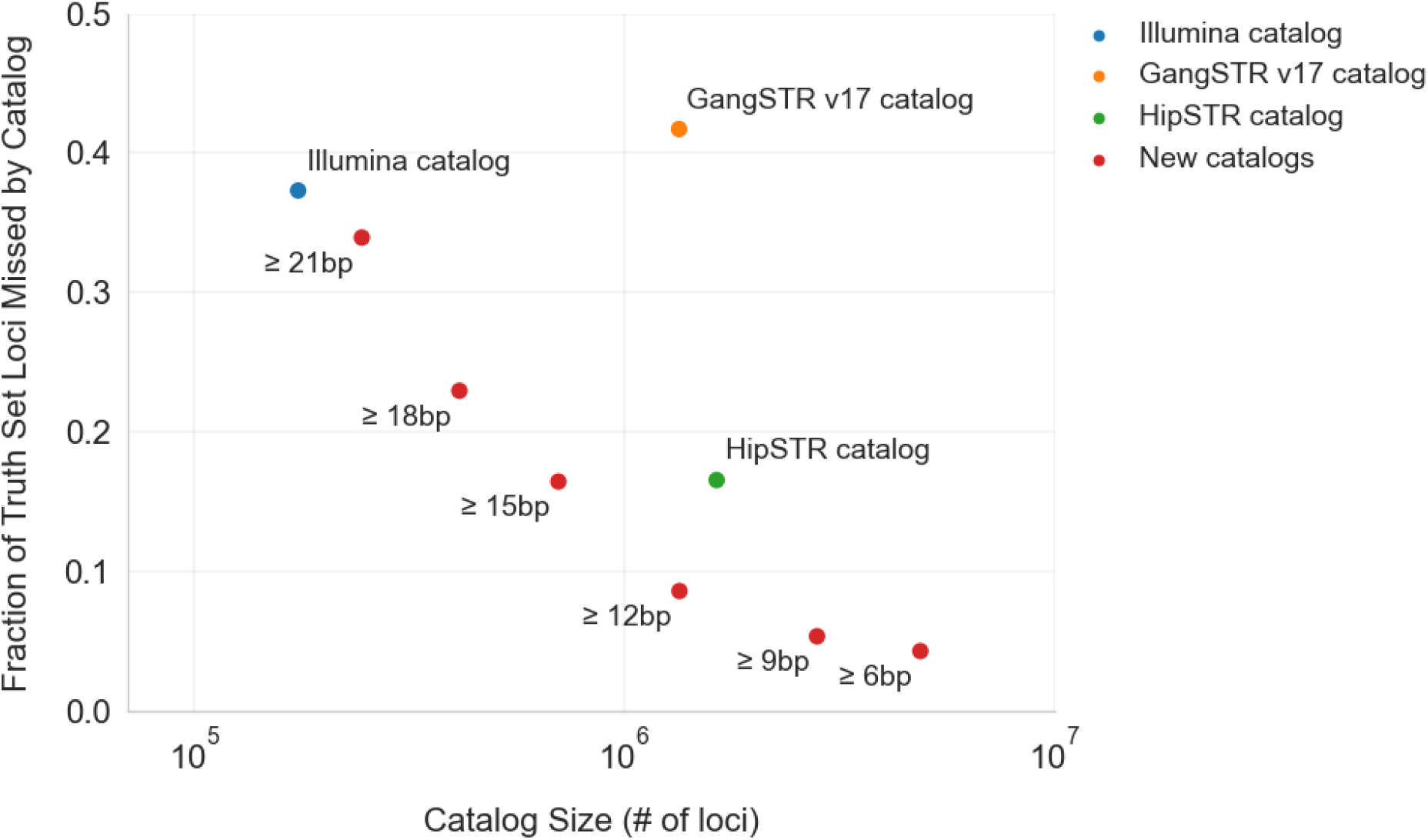
Catalogs by size and overlap with truth set loci. The scatter plot includes a dot for each of 3 publicly available TR catalogs (blue, orange, green), as well as 7 new catalogs (red) that we generated by using TRF. Each new catalog contains pure repeats in hg38 that span no less than the labeled minimum threshold of base pairs. The x axis shows the number of loci in each catalog and the y axis shows what fraction of the TR truth set is missed by this catalog. Missed loci are defined as truth set loci that don’t overlap any of the repeat loci in a catalog.

**Supplementary Figure 3:**
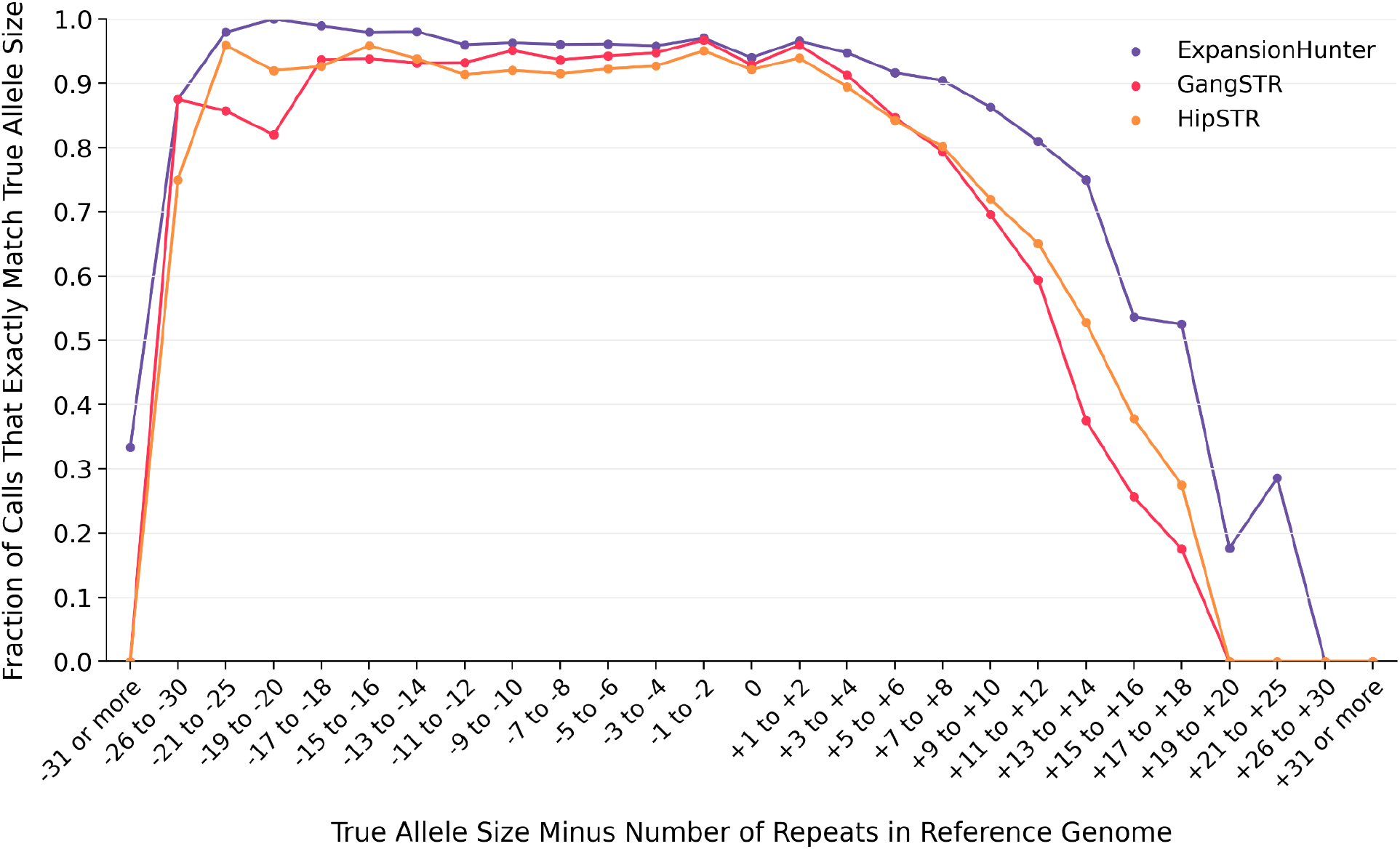
Strict accuracy after excluding no-call loci. Like Figure 4b, this plot shows the fraction of alleles that each tool got exactly right (y axis) across different true allele size bins (x axis). Here, the plot excludes all loci where one or more tools did not produce a genotype instead of treating them as equivalent to incorrect calls.

**Supplementary Figure 4:**
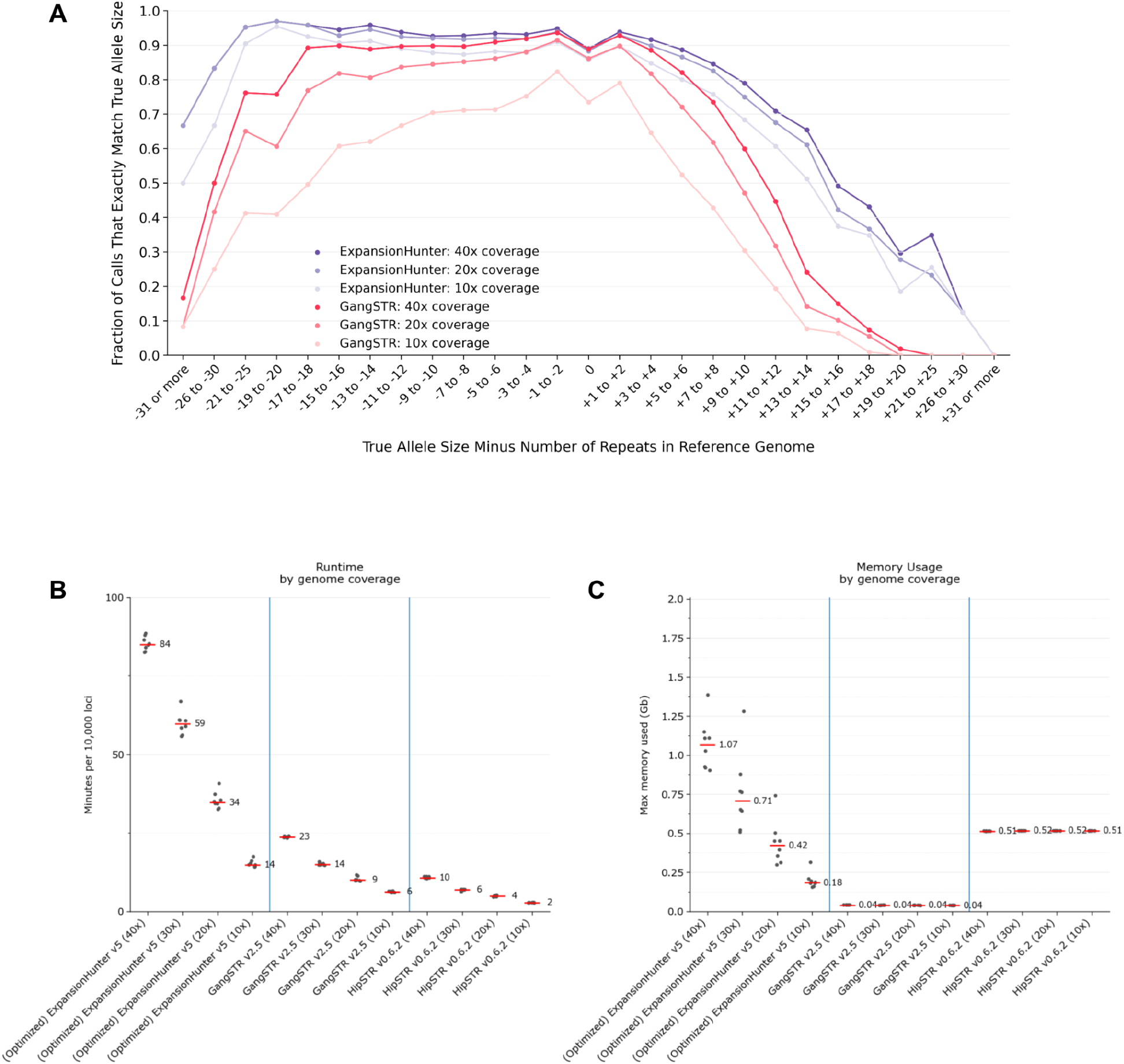
Tool accuracy, runtimes and memory use by read depth. **A. ExpansionHunter and GangSTR accuracy at different read depths.** This plot compares strict accuracy for each tool across 3 different coverage levels: 40x, 20x, 10x. **B. Tool runtimes by read depth.** For ExpansionHunter, GangSTR, and HipSTR, tool runtime in minutes per 10k loci (y-axis) is proportional to the depth of coverage of the input sample (x-axis) - as shown for coverage levels: 40x, 30x, 20x, 10x. **C. Tool memory usage by coverage.** The optimized version of ExpansionHunter uses read caching to speed up execution, and so has a higher peak memory usage (y-axis) at higher coverage levels (x-axis). Other tools’ memory usage doesn’t vary with depth of coverage.

**Supplementary Figure 5:**
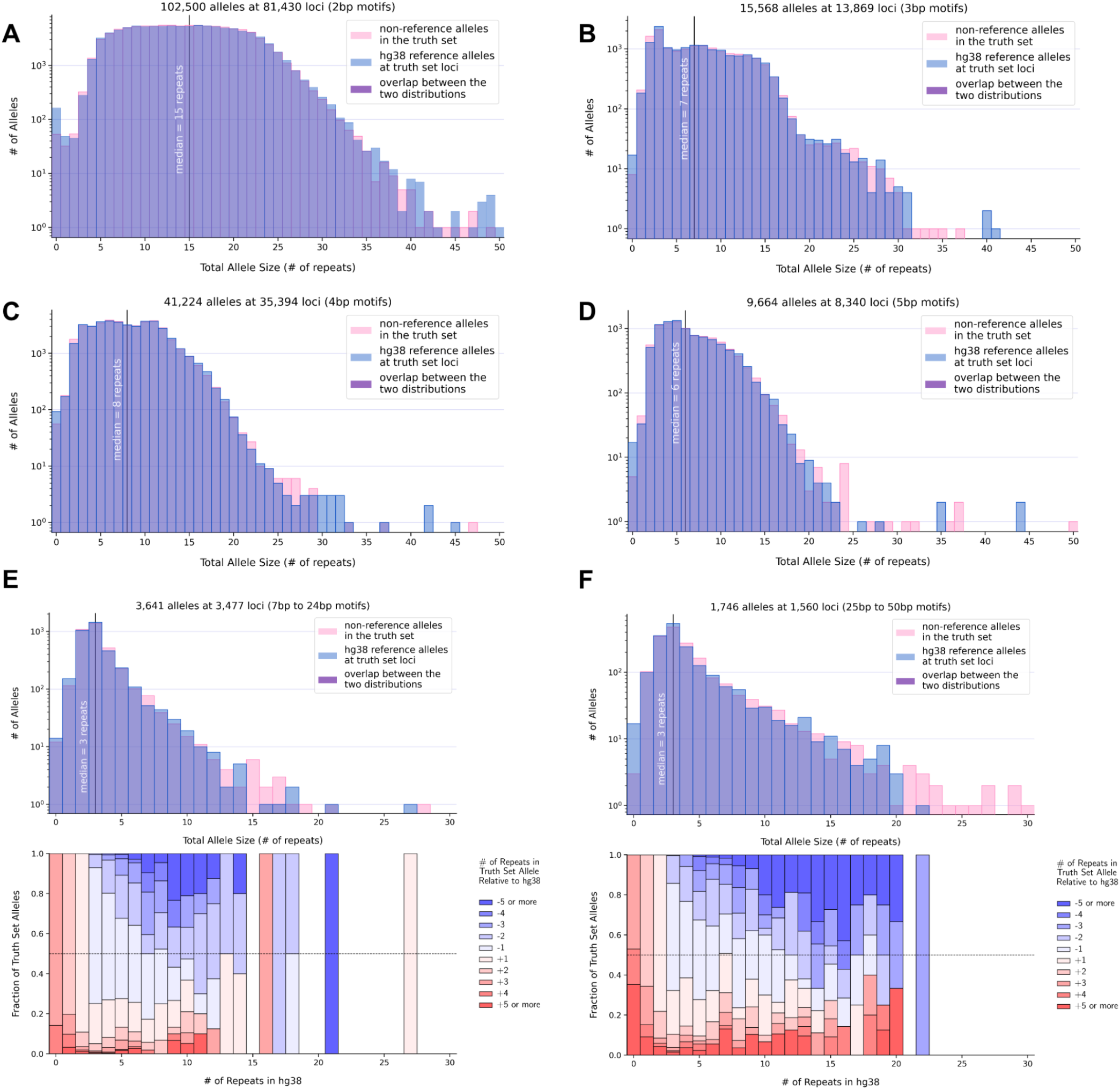
TR allele and motif sizes compared to repeat sequences in hg38 for 7-24bp and 25-50bp motifs. **A. Truth set allele sizes vs reference allele sizes with 2bp motifs.** The x axis shows allele sizes in terms of numbers of repeats. The y axis counts how many alleles of the given size (x axis) were found among truth set non-reference alleles (pink) or among hg38 reference alleles at truth set loci (blue). The two distributions - one pink, one blue - are plotted on top of each other using semi-transparent colors so that their overlap appears purple. The vertical line at x = 15 indicates the median number of repeats, which has the same value for both distributions. For the blue distribution, the reference allele size is counted twice at multi-allelic loci so that the two distributions have the same number of counts. **B. Truth set allele sizes vs reference allele sizes with 3bp motifs.** Same as panel A but for 3bp motifs. The median is at x = 7. **C. Truth set allele sizes vs reference allele sizes with 4bp motifs.** Same as panel A but for 4bp motifs. The median is at x = 8. **D. Truth set allele sizes vs reference allele sizes with 5bp motifs.** Same as panel A but for 5bp motifs. The median is at x = 6. **E. Truth set allele sizes vs reference allele sizes with 7-24bp motifs.** Same as panel A but for 7-24bp motifs. The median is at x = 3. Expansions and contractions in the truth set are shown in the lower panel using colors as in **Figure 3B**. **F. Truth set allele sizes vs reference allele sizes with 25-50bp motifs.** Same as panel A but for 25-50bp motifs. The median is at x = 3. Expansions and contractions in the truth set are shown in the lower panel using colors as in **Figure 3B**.

## Notes

https://github.com/lh3/CHM-eval/releases/tag/v0.5

https://broadinstitute.github.io/str-truth-set/html/tgg_viewer.html

https://broadinstitute.github.io/str-truth-set/html/tool_comparison_viewer.html

https://storage.googleapis.com/str-truth-set

https://github.com/broadinstitute/str-truth-set

https://github.com/broadinstitute/str-analysis

https://github.com/bw2/ExpansionHunter

https://console.cloud.google.com/storage/browser/broad-public-datasets/CHM1_CHM13_WGS2

https://console.cloud.google.com/storage/browser/broad-public-datasets/CHM1_CHM13_WES

### Competing Interest Statement

The authors have declared no competing interest.

